# NMR structures and functional roles of two related chitin-binding domains of a lytic polysaccharide monooxygenase from *Cellvibrio japonicus*

**DOI:** 10.1101/2021.04.25.441307

**Authors:** Eva Madland, Zarah Forsberg, Yong Wang, Kresten Lindorff-Larsen, Axel Niebisch, Jan Modregger, Vincent G. H. Eijsink, Finn L. Aachmann, Gaston Courtade

## Abstract

Among the extensive repertoire of carbohydrate-active enzymes, lytic polysaccharide monooxygenases (LPMOs) have a key role in recalcitrant biomass degradation. LPMOs are copper-dependent enzymes that catalyze oxidative cleavage of glycosidic bonds in polysaccharides such as cellulose and chitin. Several LPMOs contain carbohydrate-binding modules (CBMs) that are known to promote LPMO efficiency. Still, structural and functional properties of some of these CBMs remain unknown and it is not clear why some LPMOs, like *Cj*LPMO10A from *Cellvibrio* japonicus, have two CBMs (*Cj*CBM5 and *Cj*CBM73). Here, we studied substrate binding by these two CBMs to shine light on the functional variation, and determined the solution structures of both by NMR, which includes the first structure of a member of the CBM73 family. Chitin-binding experiments and molecular dynamics simulations showed that, while both CBMs bind crystalline chitin with *K*_d_ values in the µM range, *Cj*CBM73 has higher affinity than *Cj*CBM5. Furthermore, NMR titration experiments showed that *Cj*CBM5 binds soluble chitohexaose, whereas no binding to soluble chitin was detected for *Cj*CBM73. These functional differences correlated with distinctly different architectures of the substrate-binding surfaces of the two CBMs. Taken together, these results provide insight into natural variation among related chitin-binding CBMs and the possible functional implications of such variation.

## Introduction

Chitin is a linear, water insoluble polysaccharide composed of β-1,4-linked *N*-acetyl-D-glucosamine (GlcNAc) units found in the cell wall matrix of fungi and the exoskeletons of arthropods. Despite being the second most abundant polymer in nature, after cellulose, chitin does not accumulate in most ecosystems and tends to be absent in fossils (1). This is testimony to the capacity of nature to depolymerize and recycle chitin.

Chitinases (EC 3.2.1.14) catalyze the hydrolytic degradation of chitin and belong to the glycoside hydrolase (GH) class of carbohydrate-active enzymes. Even though GHs efficiently degrade amorphous regions of chitin (2–4), they are inefficient at degrading crystalline chitin (5). The discovery of lytic polysaccharide monooxygenases (LPMOs) (6, 7) has given new insights into the degradation of chitin and other structural polysaccharides. LPMOs are copper-dependent enzymes that catalyze oxidative cleavage of glycosidic bonds in crystalline polysaccharides (6, 8). Aside from chitin, LPMOs have been reported to act on polysaccharides such as cellulose (8–11), various hemicelluloses (12) and starch (13). In the degradation of chitin, LPMOs act in synergy with chitinases (4, 7). It is thought that LPMOs oxidize crystalline surfaces, causing “nicks” that lead to reduced crystallinity and introduction of new access points for chitinases (6, 10, 14).

Carbohydrate Active enZymes (CAZymes), such as chitinases and LPMOs, may just be composed of a single catalytic domain, or may contain one or more non-catalytic domains such as carbohydrate-binding modules (CBMs). Currently, the CAZy database (15) contains 88 families of CBMs with a wide variety of binding specificities, including crystalline polysaccharides and short, soluble oligosaccharides (16, 17). The major role of CBMs is to keep an enzyme in close proximity of a substrate, thereby enhancing the effective concentration of the enzyme and overall reaction efficiency (16). In the context of LPMOs, CBMs may have a particularly important role because proximity to the substrate not only contributes to enzyme efficiency, but also protects the enzyme from autocatalytic inactivation. Several studies have shown that removal of CBMs has a negative effect on LPMO performance (18–22). There are multiple families of chitin-binding and cellulose-binding CBMs, which may have different binding specificities (e.g. (18, 23)). For example, it has been shown that two cellulose-binding modules belonging to two different CBM families bind to different parts of cellulose (23). It is not trivial to predict or determine the role of CBMs and a better understanding of the ways in which they bind their substrates is needed.

To address functional variation among chitin-binding CBMs, we have used chitin-active *Cj*LPMO10A from *Cellvibro japonicus* as a model system. The catalytic domain of this LPMO, which belongs to the auxiliary activity family 10 (AA10) in CAZy, is appended to two chitin-binding CBMs, an internal family 5 CBM (*Cj*CBM5) and C-terminal family 73 CBM (*Cj*CBM73) (Figure 1). The three domains of *Cj*LPMO10A are connected by linkers that are rich in serine residues and are both approximately 30 amino acids long (Figure 1). A previous study has shown that both CBMs bind to α- and β-chitin, thus enhancing substrate-binding by the LPMO, and that the full length protein is more efficient in comparison to the catalytic domain alone (20). In the present study, we have compared multiple truncated variants of *Cj*LPMO10A (see Figure 1A) to understand the roles of the appended CBMs in LPMO functionality. Furthermore, we have used NMR spectroscopy to elucidate the solution structures of the two CBMs, *Cj*CBM5 and *Cj*CBM73, where the latter is the first structure to be determined for a member of the CBM73 family. We also used NMR titration experiments to investigate binding of the CBMs to chitohexaose. These results were complemented with molecular dynamics simulations to gain more insights into CBM binding to α-chitin. Overall, the results show that while *Cj*CBM5 and *Cj*CBM73 are similar in overall structure and both bind to crystalline chitin, they differ in apparent melting temperature, binding site architecture and the ability to bind individual chitin chains.

**Figure 1.**
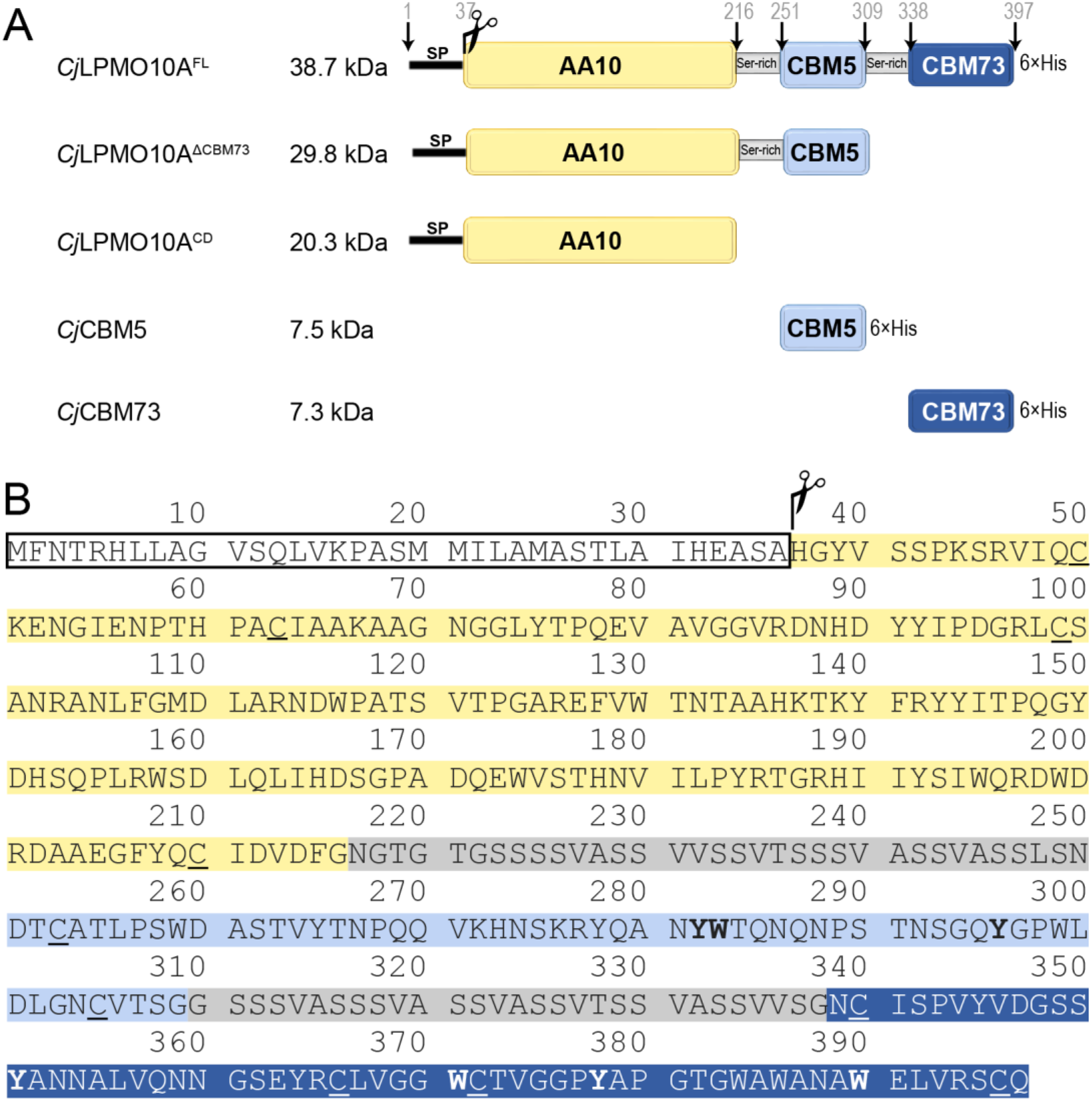
(A) Domain architecture and molecular weight of *Cj*LPMO10A and the truncated variants used in this study. The numbers above the full-length enzyme show the transitions between the domains and the linkers. The signal peptide (residue 1-37) is cleaved off during secretion. The indicated molecular weights are based on the mature protein, i.e. signal peptide free enzymes. Abbreviations used: SP, signal peptide; FL, full-length; CD, catalytic domain; Ser-rich linker; His×6, poly-histidine tag. (B) Primary sequence of *Cj*LPMO10A^FL^ with color coding according to panel A. Aromatic residues located on the binding surfaces of the two CBMs, as determined in this study, are printed in bold face; cysteine residues involved in disulfide bonds are underlined.

## Results

### The effect of CBMs on chitin oxidation is substrate concentration dependent

To better understand the functional roles of *Cj*CBM5 and *Cj*CBM73 in relation to full-length *Cj*LPMO10A we started by testing the performance of the three catalytically active versions of *Cj*LPMO10A, *Cj*LPMO10A^FL^ (FL for full-length), *Cj*LPMO10A^ΔCBM73^ (for truncation of the CBM73 domain; see Figure 1A) and fully truncated *Cj*LPMO10A^CD^ (CD for catalytic domain) at different concentrations of α-chitin (2, 10 or 50 g/L, see Figure 2). At all substrate concentrations, the full-length enzyme and the enzyme lacking only one CBM, *Cj*LPMO10A^ΔCBM73^, had similar progress curves and stayed active for the full duration of the experiment. At the two lowest substrate concentrations product formation by *Cj*LPMO10A^CD^ ceased rapidly, and faster at the lowest substrate concentration, indicative of enzyme inactivation. However, at a substrate concentration of 50 g/L all three variants showed similar progress curves and final product levels.

**Figure 2.**
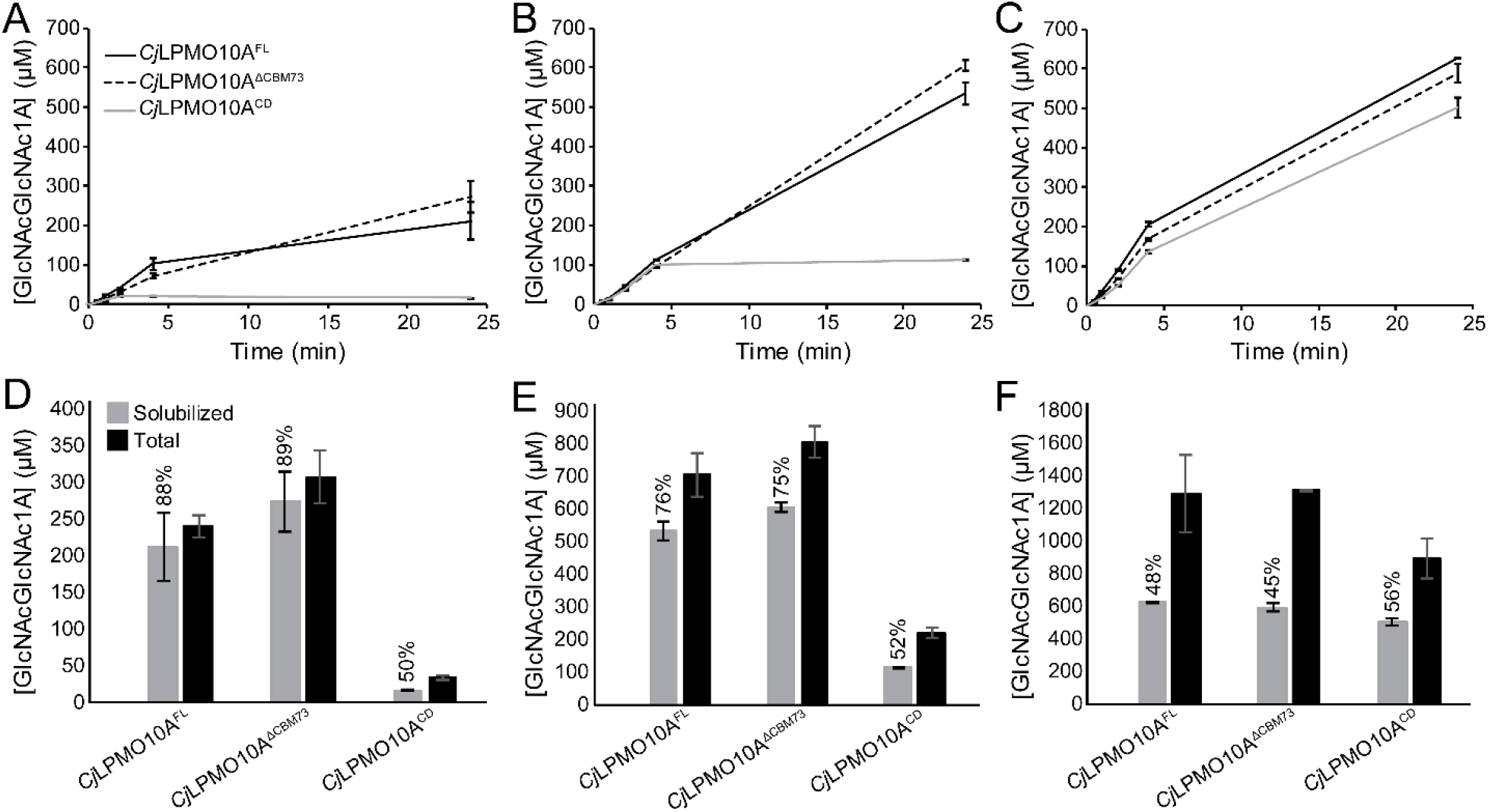
Chitin degradation by *Cj*LPMO10A variants. Panels A – C show progress curves for the formation of soluble oxidized products by *Cj*LPMO10A^FL^ (solid black line), *Cj*LPMO10A^ΔCBM73^ (dashed black line) and *Cj*LPMO10A^CD^ (solid grey line) at substrate concentrations of 2 g/L (A), 10 g/L (B) and 50 g/L (C) α-chitin. Panel D – F show quantification of solubilized (grey bars) and total oxidized sites (black bars) after 24 h of LPMO incubation at the various substrate concentrations, i.e. 2 g/L (D), 10 g/L (E) and 50 g/L (F). The fraction of soluble oxidized products is given as a percentage of the total for each reaction. All reactions were carried out with 0.5 µM LPMO and 1 mM ascorbic acid in 50 mM sodium phosphate pH 7.0 in a thermomixer set to 37 °C and 800 rpm. For quantification of soluble products, the solubilized fraction was further degraded by 0.5 µM *Sm*CHB prior to HPLC quantification. For quantification of total products (i.e. soluble and insoluble fraction), samples were heat inactivated after which all α-chitin (diluted to 2 g/L) was degraded with a combination of 2.0 µM *Sm*ChiA and 0.5 µM *Sm*CHB. The error bars show ± s.d. (n = 3).

At the two lowest substrate concentrations, the amount of soluble oxidized products (relative to the total amount) was higher for the CBM containing variants of *Cj*LPMO10A (> 85 %) compared to *Cj*LPMO10A^CD^ (about 50%) (Figure 2D-E). This indicates that, at these lower substrate concentrations, the presence of at least one CBM leads to more localized oxidation, generating a higher fraction of short soluble products, as discussed in (22) and below. At the highest substrate concentration (Figure 2F), however, the fraction of soluble oxidized products was close to 50 % for all three enzyme versions. All in all, the experiments depicted in Figure 2 did not show significant differences between the catalytic behavior of the two CBM containing variants, whereas deletion of both CBMs had a major effect.

### Thermal stability and oxidative performance

To assess possible functional differences between the full-length enzyme and the variant lacking only the CBM73, we analyzed the effect of temperature on the oxidative performance of these variants (Figure 3). It is believed that CAZymes with multiple CBMs have an advantage at elevated temperatures as the CBM(s) can counteract the loss of binding due to increased temperature (24– 26). Interestingly at the highest tested temperature (70 °C) *Cj*LPMO10A^FL^ showed significantly higher activity than *Cj*LPMO10A^ΔCBM73^. Thus, the presence of the CBM73 indeed has a beneficial effect on LPMO performance at higher temperatures. Determination of melting curves showed that the deletion of the *Cj*CBM73 had some effect on the shape of the curve but not on the apparent melting temperature of approximately 70 °C (Figure S1). The apparent melting temperatures of the isolated CBMs were 57.2°C for *Cj*CBM5 and 75.4 °C for *Cj*CBM73, whereas the apparent melting temperature of the *Cj*LPMO10A^CD^ was 70.2 °C, which was reduced to 56.6 °C upon removal of the copper.

**Figure 3.**
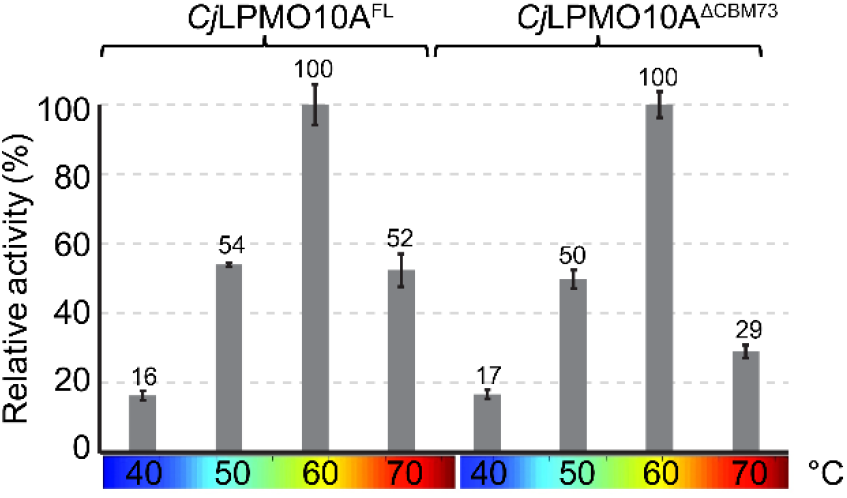
Catalytic performance of *Cj*LPMO10A^FL^ and *Cj*LPMO10A^ΔCBM73^ at varying temperatures. The relative activity was determined from linear progress curves for a 30-minute reaction. The 100 % value corresponds to 61 and 47 µM GlcNAcGlcNAc1A for *Cj*LPMO10A^FL^ and *Cj*LPMO10A^ΔCBM73^, respectively. All reactions were carried out with 0.5 µM LPMO, 10 g/L α-chitin and 1 mM ascorbic acid in 50 mM sodium phosphate pH 7.0 in a thermomixer set to the indicated temperature and 800 rpm. Prior to product quantification the solubilized fraction was further degraded with 0.5 µM *Sm*CHB. Each point represents the average of values obtained in three independent experiments.

### Solution structures of CjCBM5 and CjCBM73

The solution structures of *Cj*CBM5 (PDB: 6Z40) and *Cj*CBM73 (PDB: 6Z41) were solved by NMR spectroscopy (Figure 4, Table S1). The chemical shift assignment completion for the backbone (N, H^N^, C^α^, H^α^ and C’) and side chains (H and C) of *Cj*CBM5 (BMRB: 34519). was > 88 % and > 65%, respectively, whereas these values were > 87 % and > 59 % for *Cj*CBM73 (BMRB: 34520). Due to the cloning procedure, both proteins contained a Met at the N-terminus and an Ala followed by a 6xHis-tag at the C-terminus. For *Cj*CBM5, no resonances from these additional amino acids were assigned, whereas for *Cj*CBM73 the backbone resonances of the additional Ala and the first His in the 6xHis-tag were assigned.

**Figure 4.**
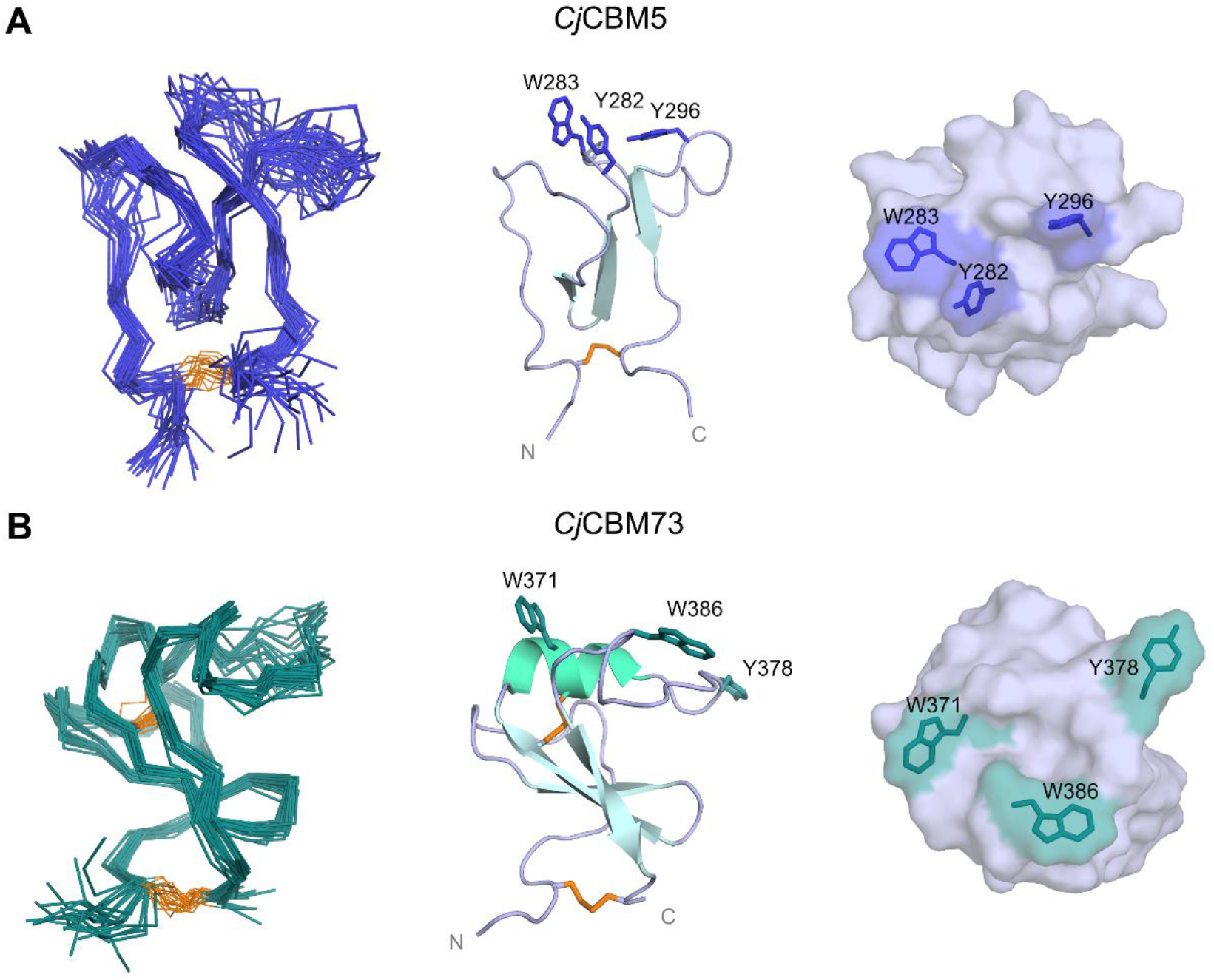
NMR solution structures of (A) *Cj*CBM5 (PDB: 6Z40) and (B) *Cj*CBM73 (PDB: 6Z41). The Figures show backbone representations for the 20 conformers with the lowest CYANA target function (left), a cartoon representation of the structure with the lowest target function (center), and a view of the binding surfaces (right). The cartoon representations also display the secondary structure elements as well as aromatic residues of the putative binding surface. Disulfide bridges (residues 253-306 in *Cj*CBM5 and 340-396 and 366-372 for *Cj*CBM73) are highlighted in orange. His-tags added for cloning purposes (see Experimental procedures) are not shown.

The structures of *Cj*CBM5 and *Cj*CBM73 are similar (C^α^ rmsd 5.6 Å) and share the same overall fold (Figure 4). This fold has previously been described (27) as an “L” shape or “ski boot” fold due to the loop region attached perpendicularly to an anti-parallel β-sheet. The structures of both CBMs are stabilized by a disulfide bridge connecting the N- and C-terminal ends of the domain. The structure of *Cj*CBM73 shows a short 3_10_ helix (residues 371-374) that is linked to the central β-strand by an additional disulfide bridge. These features are unique for the CBM73 family (see Figure S2) and lack in *Cj*CBM5 and other structurally characterized members of the CBM5 family.

Most CBMs rely on exposed aromatic residues that bind carbohydrates through CH-π interactions (16, 28). Based on structural information alone, Figure 4 shows that Y282, W283 and Y296 in *Cj*CBM5, and W371, Y378 and W386 in *Cj*CBM73 could be involved in substrate-binding. As shown in Figure S2, the aromatic pair Y282-W283 is almost fully conserved within the CBM5 family, whereas Y296 is less conserved. In the context of the CBM73 family (Figure S2), W371, Y378 and W386 appear to be highly conserved. To test interactions between chitin and these aromatic patches and their neighboring polar residues we performed NMR titrations with a soluble chitin substrate, chitohexaose, (GlcNAc)_6_.

### Probing interactions between soluble chitin and CBMs by NMR

In the case of *Cj*CBM5, titration with (GlcNAc)_6_ led to significant chemical shift perturbation for W283 and Y296 as well as for residues in the neighboring loop region (T284, Q285 and G297) that are part of the putative binding surface (Figure 5). The chemical shift perturbations were used to calculate a *K*_d_ = 2 ± 1 mM. Of note this *K*_d_ value is some three orders of magnitude higher than the value obtained with solid α-chitin (see below).

**Figure 5.**
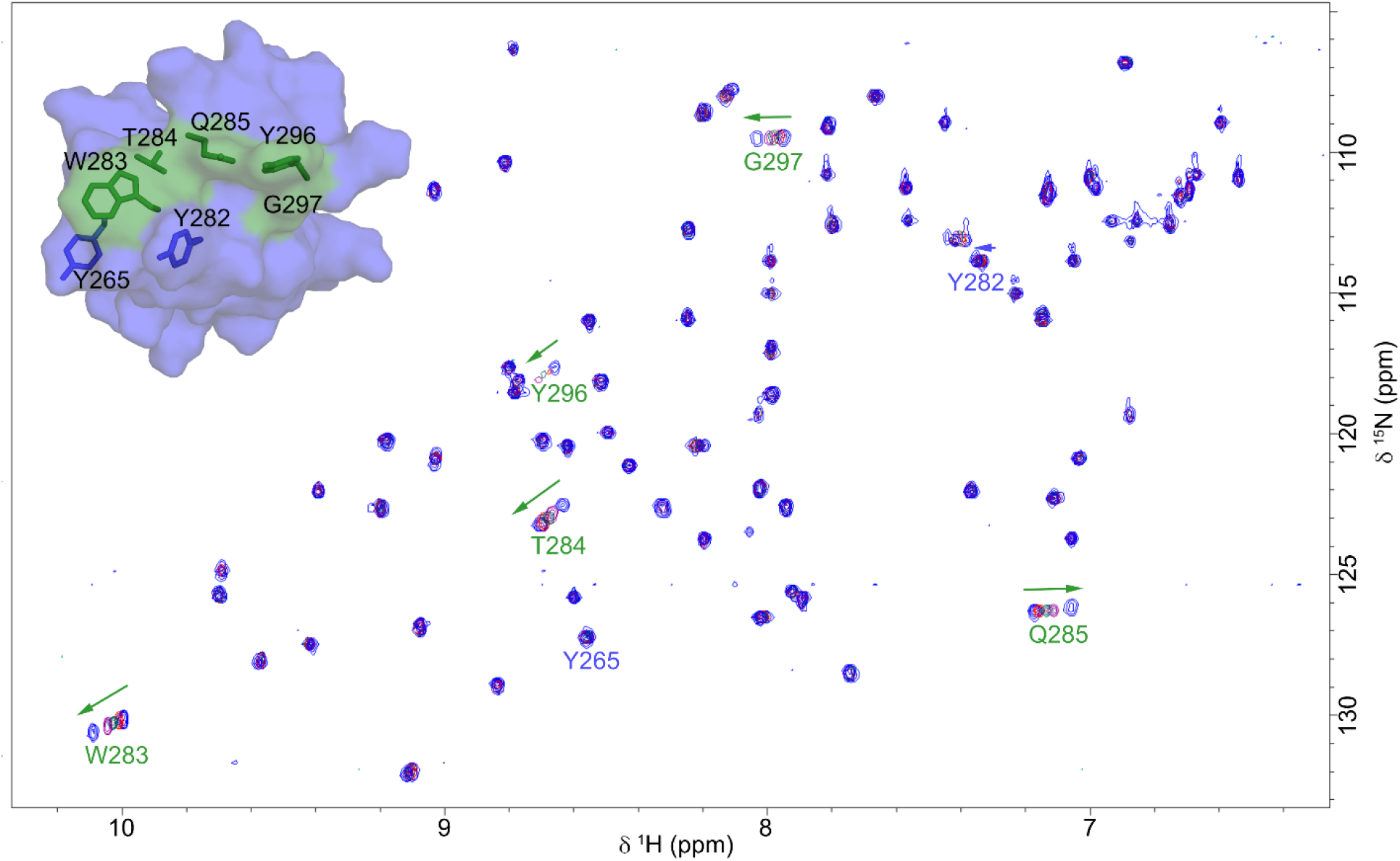
^15^N-HSQC of *Cj*CBM5 interacting with (GlcNAc)_6_. The picture shows an overlay of ^15^N-HSQC spectra for *Cj*CBM5 in the presence of (GlcNAc)_6_ at various concentrations (0.2, 1.0, 2.5 and 10 mM). The arrows indicate the direction of change in chemical shift perturbation as a result of the titration of *Cj*CBM5with (GlcNAc)_6_. Affected residues (W283, T284, Q285, Y296 and G297) are highlighted in green on the surface model of *Cj*CBM5. Other surface-exposed aromatic residues for which no significant chemical shift perturbation was detected (Y265 and Y282) are shown in blue for illustration purposes.

In stark contrast to the experiment with *Cj*CBM5, titration of *Cj*CBM73 with (GlcNAc)_6_ did not result in any significant chemical shift perturbations, indicating that this CBM does not bind this soluble substrate.

### Binding of CjCBM5 and CjCBM73 to oxidized and non-oxidized α-chitin

Previous binding studies have shown that both CBMs bind with micromolar affinity to both α- and β-chitin (20). These previous studies indicated similar *K*_d_ values (for α-chitin) for *Cj*CBM5 and *Cj*CBM73. Here we tested binding using a similar setup, using both the same batch of α-chitin and a batch of α-chitin that had been pre-oxidized with *Cj*LPMO10A^CD^ as described below and in Experimental procedures.

Oxidized chitin was prepared to assess whether surface oxidation would affect CBM binding, one idea being that gradual oxidation of the substrate surface could facilitate release of otherwise strongly bound CBMs. The material was prepared by treating chitin with the catalytic domain of *Cj*LPMO10A^CD^, followed by washing to remove solubilized oxidized chito-oligosaccharides and residual LPMO (see Experimental procedures for further details). The degree of oxidation of the solid fraction was determined upon complete enzymatic hydrolysis of the fraction, which entails that all oxidized sites end up as chitobionic acid. Data from six independent reactions, containing 20 mg/mL chitin, which corresponds to approximately 45 mM of oxidized dimer in a theoretical 100% conversion reaction, indicated a degree of oxidation of about 0.3 % (number obtained by dividing the chitobionic acid recovered from the solid fraction by the amount of chitobionic acid that would be obtained in a 100% conversion reaction). In an alternative approach, we divided the amount of chitobionic acid recovered from the solid fraction by the total amount of sugars (GlcNAc and chitobionic acid) recovered from this fraction, which indicated approximately 1% oxidation. Hence, the degree of oxidation of the insoluble fraction was estimated to be between 0.3 % and 1 % and we assume that oxidation essentially happened on the substrate surface.

Figure 6 shows binding curves for the two CBMs with “non-oxidized” (panel A and C) or “pre-oxidized” (panel B and D) α-chitin. The data show that *Cj*CBM73 (*K*_d_ = 2.9 µM) binds with slightly higher affinity than *Cj*CBM5 (*K*_d_ = 8.5 µM). The binding studies with partly oxidized chitin showed similar results. The data showed a ∼20% increase in the *K*_d_ for *Cj*CBM5, indicating that binding by this CBM may be negatively affected by surface oxidation, however, the difference was not statistically significant.

**Figure 6.**
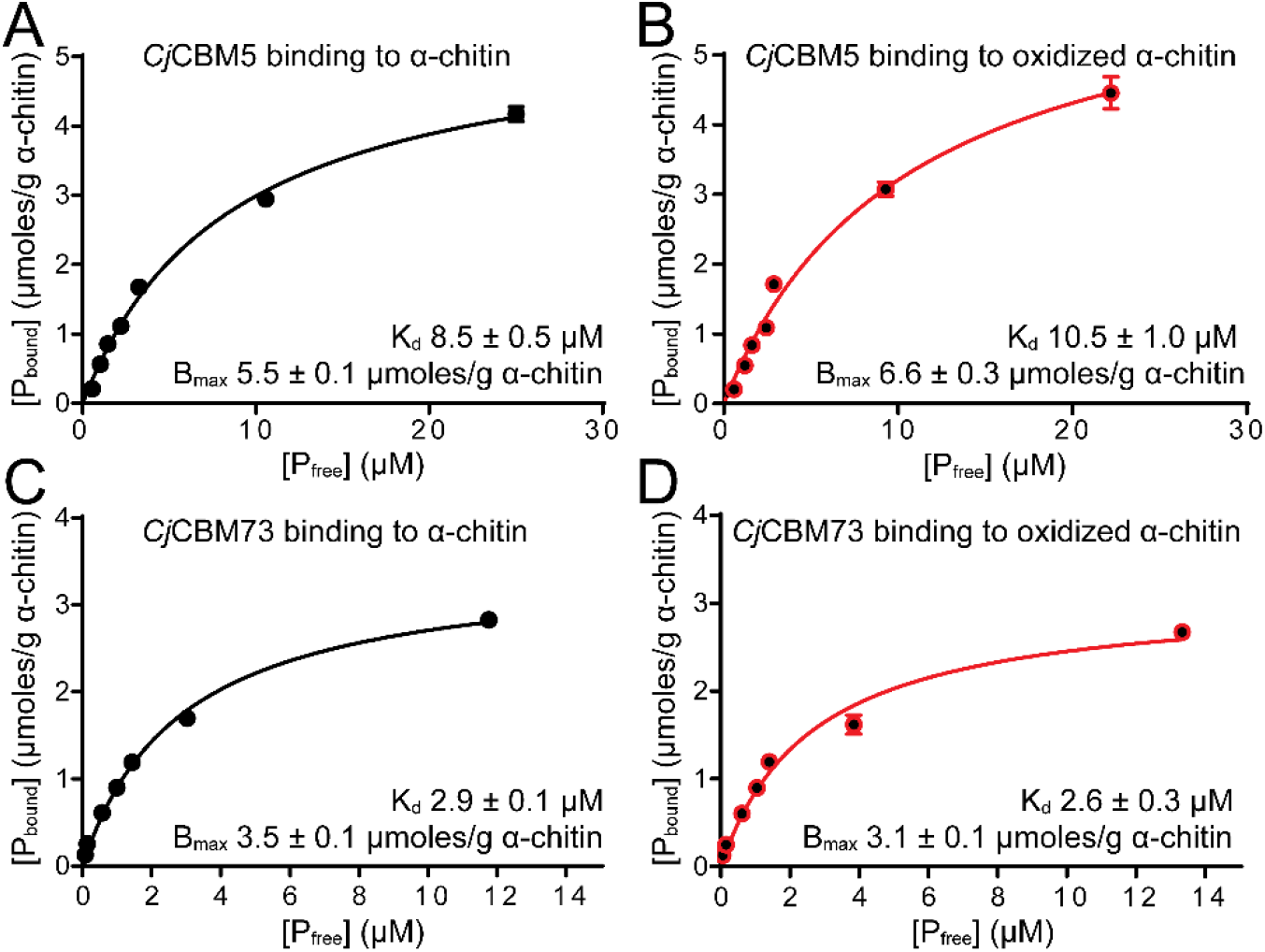
Binding of the CBMs of *Cj*LPMO10A to α-chitin. The plots show binding data for *Cj*CBM5 (A and B) and *Cj*CBM73 (C and D) incubated with α-chitin for 60 min. The experiments were carried out at 22 °C using 10 g/L α-chitin in 50 mM sodium phosphate buffer pH 7.0 and show binding of *Cj*CBM5 and *Cj*CBM73 to non-oxidized (A and C) and oxidized (B and D) substrate. P_bound_ corresponds to bound protein (μmoles/g substrate) and P_free_ corresponds to non-bound protein (μM). The error bars show ± s.d. (n = 3).

### Simulations provide insight into binding of CBMs to α-chitin

Coarse-grained (CG) simulations were performed to further investigate interactions between CBMs and a model of the surface of α-chitin. CG models based on the Martini force-field represent 3-4 atoms by a single “bead”, thereby reducing the number of particles that are simulated (29). This allows simulations to be run longer and to sample longer time scales, compared to atomistic simulations. We combined CG models of chitin and the CBMs with well-tempered metadynamics (WT-MetaD) simulations to further enhance sampling of CBM-chitin binding/unbinding events, which occur on long time scales. In the WT-MetaD approach, protein conformations along a set of collective variables are biased by a history-dependent potential. The total bias (i.e. sum of the Gaussians in the potential) forces the system to escape from local free-energy minima and explore different regions of the collective variable space. In the case of the CBM-chitin model we used two collective variables as proxies for binding: (i) the Euclidean distance (*r*_*chitin*_) and (ii) number of contacts, ⟨c_w_⟩, between aromatic residues in the putative substrate binding surfaces (*Cj*CBM5: Y282, W283, Y296; *Cj*CBM73: W371, Y378, W386) and the chitin surface. Details on the calculation of these collective variables are provided in the Experimental procedures.

To promote binding to chitin by the CBMs it was necessary to rescale the interaction strengths between chitin beads and protein beads in the Martini model (see Experimental procedures for details). The effect of rescaling these interactions by 0% (unchanged) or by an up to 15% increase in the strength of the chitin-protein interaction was evaluated by running umbrella-sampling simulations (Figure S3) on the rescaled models, and by comparing dissociation constants calculated from these simulations with experimentally determined values (Figure 6). The results (Table 1) show that the best agreement with experiment was attained with a 10% increase in the chitin-protein interaction strength. The free-energy surfaces of *Cj*CBM5 and *Cj*CBM73 have similar appearances, but *Cj*CBM73 has a deeper well than *Cj*CBM5, which correlates with its experimentally observed stronger affinity for chitin (Figure S3).

**Table 1.** Dissociation constants (*K*_d_) for binding of the *Cj*CBMs to α-chitin, determined by experiments (see Figure 6) and simulations (see Figures 7 and S3). The modeled values were calculated from umbrella sampling simulations in which the interaction strength between chitin beads and protein beads remained unchanged (0%) or was increased by 5%, 10% and 15%.

| Protein | Simulations ($\mu\text{M}$) | | | | Experiments ($\mu\text{M}$) |
| --- | --- | --- | --- | --- | --- |
|  | 0% | 5% | 10% | 15% |  |
| *Cj*CBM5 | 1800 | 480 | 36 | 0.65 | $8.9 \pm 0.5$ |
| *Cj*CBM73 | 290 | 76 | 15 | 0.13 | $2.9 \pm 0.1$ |

The number of contacts between all amino acids in each CBM and the α-chitin surface was calculated for every frame (n=10000) in the WT-MetaD simulation and reweighted using the bias from the simulation (see Experimental procedures for details). The results (Figure 7) show which residues have the most contacts, i.e. ⟨c_w_⟩ > 0.5, with the substrate over time. In the case of *Cj*CBM5 (Figure 7A,C), regions with most contacts include, and are to a large extend limited to, the three aromatic residues of the putative binding surface (Y282, W283, Y296). Additionally, the region around Y265 seems to be somewhat involved in substrate binding albeit with much fewer contacts. These observations are in good agreement with the chemical shift perturbation data for binding of (GlcNAc)_6_. Similar observations were made for *Cj*CBM73 (Figure 7B,D), in the sense that also in this case the interacting regions include, and are to a large extend limited to, the three aromatic residues of the putative binding surface (W371, Y378, W386). Furthermore, also in this case interactions with fewer contacts (0.2 < ⟨c_w_⟩ < 0.5) with a fourth aromatic residue, Y351, were observed. In order to match the experiments as closely as possible, we included the C-terminal His in the simulations and found that these have a number of contacts with the substrate. (Figure 7A,B). All in all, these analyses show that the amino acids on the surface of *Cj*CBM5 that most frequently interact with chitin form a relatively linear arrangement (Figure 7E), perhaps reflecting that interactions are limited to a single chitin chain, whereas the arrangement of aromatic amino acids on the surface of *Cj*CBM73 is wider and suggests a more extended substrate-binding surface (Figure 7F).

**Figure 7.**
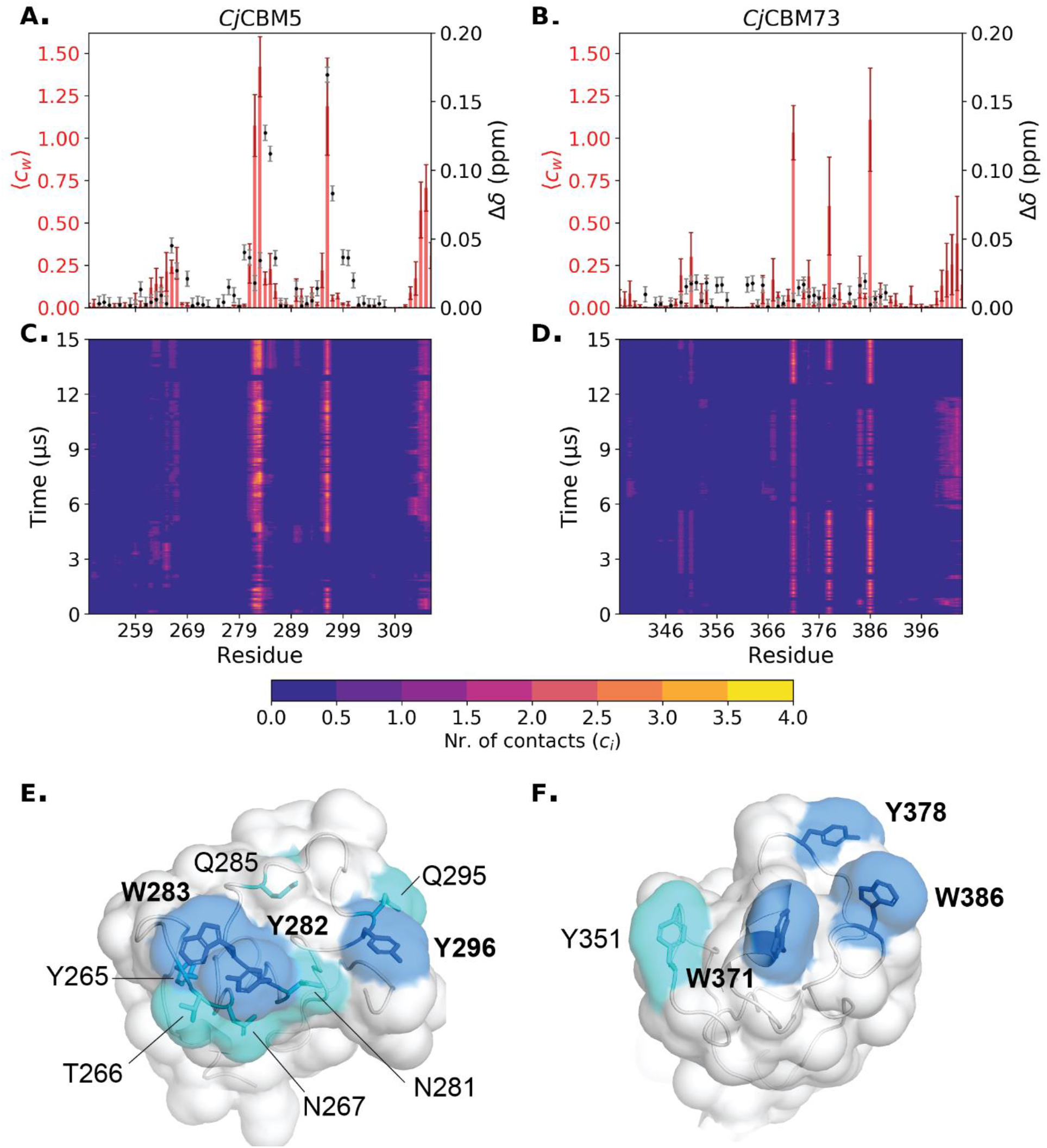
Chitin binding probed by NMR and simulations for *Cj*CBM5 and *Cj*CBM73. Panels A and B show the weighted average number of contacts observed during the simulation (⟨*c*⟩_*w*_; red bars with dark red error bars) and the chemical shift perturbations (Δδ; black dots with grey error bars) observed by NMR upon addition of (GlcNAc)_6_. The error bars for the number of contacts were calculated using block analysis (30); error bars for chemical shift perturbations correspond to 0.003 ppm. Note that no significant chemical shift perturbations were recorded for *Cj*CBM73. Panels C and D show the number of contacts between each amino acid and the α-chitin surface per frame of the 15 µs simulations, *c*_*i*_, using a cut-off distance of 0.3 nm (see Experimental procedures for details). We note that due to the use of coarse-grained models and because of the use of metadynamics, i.e. enhanced sampling, the timescales do not here correspond to a physical time scale. Panels E and F show the substrate-binding surfaces of representative conformations of the bound state of *Cj*CBM5 and *Cj*CBM73, respectively. The side chains of amino acids on the binding surface that have most contacts (⟨*c*_*w*_⟩ > 0.5) with chitin are colored blue, while the side chains of amino acids with fewer contacts (0.2 < ⟨*c*_*w*_ ⟩ < 0.5) are colored cyan.

## Discussion

In multi-modular LPMOs, CBMs tethered to the LPMO domain have significant impact on the catalytic efficiency of the enzyme (19, 20, 22, 31). Therefore, it is important to gain a deeper understanding of the mechanisms by which CBMs recognize and bind their target substrates. Here, we have investigated two CBMs from *Cj*LPMO10A, *Cj*CBM5 and *Cj*CBM73, to illuminate structural and functional differences between these chitin-binding domains. The present results include the first structure for a member of the CBM73 family.

The NMR solution structures show that, although both CBMs have similar overall folds, *Cj*CBM73 has a 3_10_-helix connected by an additional disulfide bridge. These features appear to be conserved in CBM73 family (Figure S2). To obtain further insight into the structural variation between small chitin-binding CBMs, we compared the structures of *Cj*CBM5 and *Cj*CBM73 with the structures of five CBM5s and a CBM12 (Figure 8). The CBM12 is included since the CBMs in this family are closely related to family 5 CBMs (16). It has previously been shown (32–34) that, in addition to conserved surface-exposed aromatic residues, these CBMs share two additionally conserved aromatic amino acids (Y265 and W299 in *Cj*CBM) that also occur in CBM73s (Y351 and W390 in *Cj*CBM73; Figure S2). These residues are a part of the hydrophobic core of the proteins. All CBMs (32–38) in Figure 8 bind chitin.

**Figure 8.**
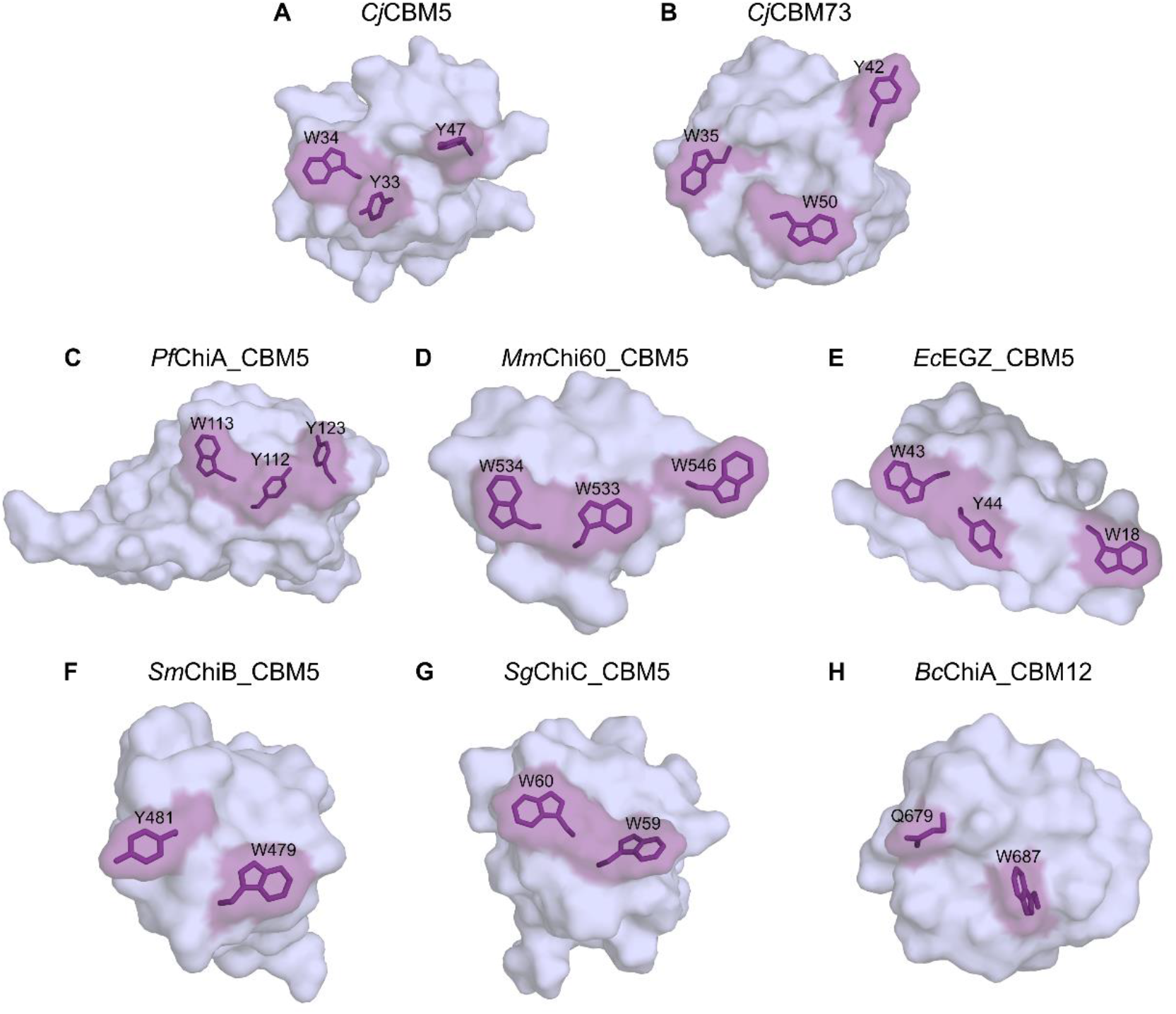
Comparison of the binding surfaces of the NMR structures of *Cj*CBM5 and *Cj*CBM73 with the structures of other CBM5 domains and one CBM12 domain. The other structures are derived from: *Pf*ChiA_CBM5 (34) (PDB ID: 2RTS; NMR structure), *Mm*Chi60_CBM5 (37) (PDB ID: 4HMC; X-ray diffraction structure), *Ec*EGZ_CBM5 (27) (PDB ID: 1AIW; NMR structure), *Sm*ChiB_CBM5 (36) (PDB ID: 1E15; X-ray diffraction structure), *Sg*ChiC_CBM5 (33) (PDB ID: 2D49; NMR structure) and *Bc*ChiA_CBM12 (32, 38) (PDB ID: 1ED7; NMR structure). Residues shown or predicted to be involved in substrate binding are highlighted in purple.

Previous studies (33, 34) have established the importance of the two consecutive, conserved aromatic residues, Y-W or W-W, in family 5 CBMs (Y282-W283 in *Cj*CBM5). Site-directed-mutagenesis studies have shown that a third aromatic residue, Y296, present on the surface of *Cj*CBM5, *Pf*ChiA_CBM5 and *Mm*Chi60_CBM5, also contributes to chitin-binding (34). Whereas the NMR titration experiment with soluble chitohexaose did not show binding for *Cj*CBM73, results for *Cj*CBM5 showed that both W283 and Y296 are involved in binding (GlcNAc)_6_. Additionally, the polar residues T284 and Q285 also appear to contribute to binding (GlcNAc)_6_. These observations suggest that chitin-binding by *Cj*CBM5 likely involves a combination of CH–π interactions (28) and hydrogen bonding. This is in good agreement with previous observations by Akagi et al. (33) who studied binding of to (GlcNAc)_6_ to *Sg*ChiC_CBM5. Like *Cj*CBM5 (*K*_d_ = 2 ± 1 mM), *Sg*ChiC_CBM5 binds (GlcNAc)_6_ with low mM affinity (*K*_d_ = 1.6 ± 0.3 mM).

*Cj*CBM5, like other CBM5s in Figure 8, has three exposed aromatic residues with a close to linear arrangement of the side chains on the surface. This type of arrangement is often found in cellulose-binding domains (39–41), where the distance between the three aromatic residues coincides with the spacing of every second glucose ring in a single chain (40, 42). Compared to the other CBMs in Figure 8, the arrangement of the three exposed aromatic residues in *Cj*CBM73; W371, Y378 and W386, differs, which suggests that *Cj*CBM73 has a wider binding surface that may interact with several chitin chains. This could explain why *Cj*CBM73 cannot bind (GlcNAc)_6_, a single-chain analog, while *Cj*CBM5 can.

The distinct arrangements of amino acids on the binding surfaces of *Cj*CBM5 and *Cj*CBM73 may also explain the experimentally and computationally observed differences in binding to α-chitin (Figures 6 and 7). The side chains of the amino acids with most chitin contacts in the simulations (Figure 7E) form a linear arrangement in *Cj*CBM5 but are distributed on a larger and wider surface in *Cj*CBM73 (Figure 7F). Both experiment and simulations indicated that CjCBM5 binds to chitin with lower affinity compared to *Cj*CBM73 (Table 1), whereas this is the other way around for (single chain) (GlcNAc)_6_. The stronger affinity of *Cj*CBM73 for insoluble α-chitin can be explained by its binding surface covering a larger area than the binding surface of *Cj*CBM5.

At low substrate concentrations, the catalytic performance of *Cj*LPMO10A^FL^ and *Cj*LPMO10A^ΔCBM73^ is superior to that of *Cj*LPMO10A^CD^ and the progress curves in Figure 2A-B show that this is due to rapid inactivation of *Cj*LPMO10A^CD^. Forsberg et al (20) have previously shown that almost all the binding affinity for chitin in *Cj*LPMO10A^FL^ resides on the CBMs. The strong binding provided by the CBMs ensures that the LPMO stays close to its substrate, thus increasing the chances that the interaction of the reduced catalytic domain with the oxygen co-substrate leads to a productive reaction (i.e., cleavage of chitin) rather than futile turnover that may lead to auto-catalytic enzyme inactivation (43), as has previously been observed for other CBM-containing LPMOs (22, 31). At the highest substrate concentration (Figure 2C), the efficiency of all enzyme variants was approximately the same, likely because the high substrate load favors *Cj*LPMO10A^CD^ binding to chitin, reducing the frequency of futile turnovers and the concurrent risk of enzyme inactivation. This observation is in agreement with a previous study (22) showing that the negative effect of truncation of the CBM2 from a two-domain cellulose-active LPMO was smaller at higher substrate concentrations.

The protective effect of the substrate, mediated by the CBMs, became more evident at higher temperatures (Figure 3), where *Cj*LPMO10A^FL^ showed higher catalytic performance than *Cj*LPMO10A^ΔCBM73^, indicating that *Cj*CBM73 appears to provide additional protection to the enzyme from thermal inactivation. It is conceivable that the increased performance at higher temperatures translates into increased performance at lower, more physiologically relevant temperatures where the enzyme may experience other types of stress, such as very low substrate concentrations or high levels of oxidant.

The abovementioned previous study with a two-domain cellulose LPMO (22) shows that the anchoring effect of the CBMs leads to a higher fraction of soluble oxidized products relative to oxidized sites on the insoluble substrate. A similar effect was also observed for *Cj*LPMO10A^FL^ and *Cj*LPMO10A^ΔCBM73^ which produced a higher fraction of soluble oxidized products compared to *Cj*LPMO10A^CD^ binding (Figure 2D-F). Interestingly, this difference became less at higher substrate concentrations, which is likely due to the fact that higher substrate concentrations increase the chance that a substrate-anchored but otherwise freely moving (22) catalytic domain acts on a neighboring fibril rather than the fibril to which it is bound.

Considering the LPMO reaction cycle and considering that anchoring by the CBMs could lead to multiple oxidized sites localized on the chitin surface around the CBM binding site, it is conceivable that accumulation of oxidized sites could trigger unbinding of the CBMs. The results of our attempts to test this hypothesis by studying CBM binding to partially oxidized chitin (Figure 6) were not conclusive, but did indicate that substrate oxidation slightly weakened chitin binding by *Cj*CBM5.

In conclusion, we have revealed structural and functional variation between the two chitin-binding domains in *Cj*LPMO10A. While it is clear that the presence of these CBMs has a significant effect on the catalytic performance of the LPMO, the reason for why nature has evolved enzymes with two slightly different chitin-binding domains remains unclear. It is conceivable that the combination of these domains provides advantages when acting on natural chitin-rich substrates, which likely are complex co-polymeric structures that may show structural variations. It is possible that functional differences between the CBM5 and the CBM73 remain undetected in the present experiments with a rather homogeneous, heavily processed chitin. Eventually, insights into these two CBMs will increase our understanding of how LPMOs depolymerize insoluble polysaccharides.

## Experimental procedures

### Cloning, expression and purification of CjLPMO10A variants

The gene encoding *Cj*LPMO10A^FL^ (residue 1-397) was codon optimized for *E. coli* expression. *Cj*LPMO10A^CD^ (residue 1-216) was cloned into the pRSET B expression vector (Invitrogen) as previously described (20), as well as the construct lacking the CBM73 and the preceding poly-serine linker, named *Cj*LPMO10A^ΔCBM73^ (residue 1-307).

To obtain better expression of *Cj*LPMO10A^FL^, the codon optimized gene encoding mature *Cj*LPMO10A^FL^ (residues 37-397) was cloned behind an IPTG-inducible T5 promoter in the pD441-CH expression vector by ATUM (Newark, CA, USA), resulting in a fusion construct with an N-terminal *E. coli* OmpA signal peptide and a C-terminal His_6_ motif (Gly-(His)_6_). The expression vector was transformed into chemically competent *E. coli* BL21 (New England Biolabs). Production of *Cj*LPMO10A^FL^ was achieved by fed-batch fermentation of the expression strain in a 1-liter fermenter (DASGIP® benchtop bioreactors for cell culture; Eppendorf, Hamburg, Germany), essentially as described previously (44), with the following modifications: at the start of the feed phase, the temperature was switched to 25 °C, and 0.6 mM isopropyl-β-D-thiogalactopyranoside (IPTG) was added to the glucose feed solution for continuous induction of gene expression. After 18 h of glucose feed, the cells were removed by centrifugation. The culture supernatant containing the target protein was concentrated 3-fold and buffer exchanged against 6 volumes of working buffer (50 mM Tris/HCl, 300 mM NaCl, pH 8.0) by crossflow filtration (Millipore Pellicon 2 mini filter, regenerated cellulose, 3 kDa MWCO). After centrifugation for 30 min at 35,000 x g to remove precipitated proteins and filtration through a 0.2 µm Nalgene Rapid-Flow sterile bottle-top filter unit (Thermo Scientific, Waltham, MA, USA), the culture filtrate was applied to a 20-mL nickel–nitrilotriacetic acid–sepharose column connected to an ÄKTA express FPLC system (GE Healthcare Lifesciences). After washing with 10 column volumes of working buffer containing 20 mM imidazole, bound protein was eluted with a buffer containing 200 mM imidazole. Fractions containing the target protein were pooled and buffer exchanged into 20 mM Tris/HCl, 200 mM NaCl, pH 7.5 by gel filtration over Sephadex G25 (GE Healthcare, 4 x HiPrep Desalting 26/10 columns).

*Cj*LPMO10A^ΔCBM73^ and *Cj*LPMO10A^CD^ were expressed in Lysogeny broth (LB) media containing 50 μg/mL ampicillin. Cells harboring the plasmid was grown at 30 °C for 24 h, without any induction, prior to harvest. The protein was extracted from the periplasmic space using an osmotic shock method that was first described by Manoil & Beckwith (45), followed by purification using a two-step chromatography protocol. The periplasmic extract was adjusted to 50 mM Tris/HCl pH 9.0 (loading buffer) and loaded onto a 5 mL Q Sepharose anion exchange column (GE Healthcare). Proteins were eluted using a linear salt gradient (0-500 mM NaCl) over 60 column volumes using a flow rate of 2.5 mL/min. LPMO containing fractions were pooled and concentrated to 1 mL before being loaded onto a HiLoad 16/60 Superdex 75 size exclusion column (GE Healthcare) operated with a running buffer consisting of 50 mM Tris pH 7.5 and 200 mM NaCl, at a flow rate of 1 mL/min. Fractions containing pure LPMO were identified by SDS-PAGE and subsequently pooled and concentrated using Amicon Ultra centrifugal filters (Millipore) with a molecular weight cut-off of 10 kDa. Protein concentrations were measured using the Bradford assay (Bio-Rad). The protein solutions were stored at 4 °C until further use.

Expression plasmids for *Cj*CBM5 (residue 251-309) and *Cj*CBM73 (residue 338-397) based on the pNIC-CH vector (Addgene) were used for cytoplasmic expression as previously described (20). This cloning procedure adds a Met residue to the N-terminus as well as one Ala residue and a poly-histidine tag (6xHis-tag) to the C-terminus of both proteins. Pre-cultures in 5 mL LB-medium (10 g/L tryptone, 5 g/L yeast extract and 5 g/L NaCl) were used to inoculate 500 mL of TB-medium (Terrific Broth) supplemented with 50 µg/mL kanamycin. The cultures were grown at 37 °C for approximately 3 hours in a LEX-24 Bioreactor (Harbinger Biotechnology, Canada) using compressed air for aeration and mixing. Expression was induced by adding IPTG to a final concentration of 0.1 mM at an optical density at 600 nm (OD_600_) of 0.6, followed by incubation for 24 h at 23°C. Cells were harvested by centrifugation (5,500 × g, 10 min) followed by cell lysis using pulsed sonication in a buffer containing 50 mM Tris/HCl pH 8.0, 500 mM NaCl and 5 mM imidazole. Cell debris was removed by centrifugation (75,000 × g, 30 min) and the supernatant was loaded onto a 5 mL HisTrap HP Ni Sepharose column (GE Healthcare) equilibrated with lysis buffer. The protein was eluted by applying a 25 CV linear gradient to reach 100 % of a buffer containing 50 mM Tris/HCl pH 8.0, 500 mM NaCl and 500 mM imidazole, at a flow rate of 2.5 mL/min. Protein containing fractions were analyzed by SDS-PAGE and subsequently concentrated, with concomitant buffer exchange to 20 mM Tris/HCl pH 8.0, using an Amicon® Ultra centrifugal filter (Millipore) with 3 kDa cut-off. The concentrations of *Cj*CBM5 and *Cj*CBM73 were determined by measuring A_280_ and calculated using theoretical molar extinction coefficients (ε_280, *Cj*CBM5_ = 22,585 M^-1^cm^-1^, ε_280, *Cj*CBM73_ = 28,210 M^-1^cm^-1^).

### Production of CjCBM5 and CjCBM73 for NMR studies

*Cj*CBM5 and *Cj*CBM73 samples for NMR studies were produced both with ^13^C and ^15^N isotopic labeling and ^15^N labeling only. A preculture was grown in 6 mL LB medium supplemented with 50 μg/mL kanamycin in a shaking incubator at 30 °C, 225 rpm, for six hours. A main culture of 500 mL M9 medium (6 g/L Na_2_HPO_4_, 3 g/L KH_2_PO_4_, 0.5 g/L NaCl) supplemented with 500 μg/mL kanamycin, 0.5 g (^15^NH_4_)_2_SO_4_, 6 mL glycerol, 5 mL ^15^N Bioexpress Cell Growth Medium (Cambridge Isotope Laboratories, Tewksbury, MA, USA), 5 mL of Gibco MEM Vitamin Solution (100x), 1 mL MgSO_4_ (1 M) and 5 mL of a trace-metal solution (0.1 g/L ZnSO_4_, 0.8 g/L MnSO_4_, 0.5 g/L FeSO_4_, 0.1 g/L CuSO_4_, 1 g/L CaCl_2_) was inoculated with 1 % of the preculture and incubated at 22 °C in a LEX-24 Bioreactor as described above. After 18 hours, the cultures were induced with 0.5 mM IPTG to a final concentration of 0.5 mM was followed by incubation at 22°C for 24 h. Cells were harvested by centrifugation at 4 °C, 6000 × g, for 5 min. The pellet was resuspended in 20 mL lysis buffer (50 mM Tris-HCl, 50 mM NaCl, 0.05 % Triton X-100, pH 8.0) supplemented with 1 tablet EDTA-free cOmplete ULTRA protease inhibitor (Roche) followed by pulsed sonication. Cell debris was removed by centrifugation at 4 °C, 16,600 × g, for 45 min. The supernatant was sterilized by filtration through a 0.2 µm Sterile-flip filter unit (Nalgene). Buffer B (50 mM Tris-HCl, 400 mM imidazole, pH 8.0) was added to the filtered lysate to obtain a final concentration of 20 mM imidazole. The proteins were purified by loading the supernatant onto a 1 mL HisTrap HP Ni-sepharose column (GE Healthcare Life Sciences) equilibrated with 5 CV of 95% Buffer A (50 mM Tris-HCl, pH 8.0) and 5 % Buffer B with a flow rate of 1 mL/min. Impurities were removed by washing with 95% Buffer A and 5 % Buffer B for 10 CV. The protein was eluted using a 30 CV gradient of 5-100% Buffer B. The purity of the protein fractions was assessed with SDS-PAGE.

The protocol for production and purification of non-labeled samples of *Cj*CBM5 and *Cj*CBM73 for NMR studies was done as described above, except that 2xLB medium (20 g/L tryptone, 10 g/L yeast extract and 5 g/L NaCl) was used instead of M9.

Fractions shown to contain *Cj*CBM73 were pooled and concentrated using Amicon Ultra protein concentrators (MWCO 3 kDa) at 10 °C and 7000 × g to obtain a volume of ∼5 mL. This protein solution was loaded onto a size-exclusion chromatography (SEC) column (HiLoad 16/600 Superdex 75 pg; 120 mL CV) that had been equilibrated with SEC-buffer pH 7.5 (50 mM Tris-HCl, 20 mM NaCl) for 1 CV. Protein fractions were eluted using a 1 mL/min flow rate and the concentration was measured as mentioned above.

The buffer in the protein-containing fractions was exchanged to NMR buffers (for structure elucidation: 25 mM sodium phosphate and 10 mM NaCl, pH 5.5; for interaction studies: 50 mM sodium phosphate (*Cj*CBM5) or 25 mM sodium phosphate (*Cj*CBM73), pH 7.0) prior to concentrating to ∼70 µM and a final volume of ∼400 µL. All steps were performed by centrifugation using Amicon Ultra protein concentration tubes (MWCO 3 kDa) at 10 °C and 7000 × g. NMR samples were prepared by adding D_2_O to a final ratio of 90 % H_2_O/10% D_2_O.

### Chitin degradation experiments

Unless stated otherwise, reactions were performed with 0.5 µM LPMO in 50 mM sodium phosphate buffer pH 7.0 in the presence of 1 mM ascorbic acid at 37 °C and 800 rpm in an Eppendorf thermomixer. All reactions were performed in triplicates.

### Preparation of oxidized chitin for binding studies

*Cj*LPMO10A^CD^ which is known to bind weakly to α-chitin (20) and which was expected to oxidize the chitin surface more randomly compared to the full length enzyme (Figure 1; (22)) was used to prepare oxidized chitin. Six 1 mL reactions, each containing 20 g/L α-chitin suspended in 50 mM sodium phosphate pH 7.0, were supplemented with 1 µM *Cj*LPMO10A^CD^ and 1 mM ascorbic acid three times with 24 h intervals (i.e. to a final concentration of 3 µM enzyme and 3 mM ascorbic acid). The reactions were incubated in a thermomixer set to 37 °C and 800 rpm. After 72 h of incubation, samples were taken from all six reactions and diluted in buffer supplemented with 2 µM *Sm*ChiA (46) and 0.5 µM *Sm*CHB (47) to a substrate concentration of 2 g/L. These reaction mixtures were incubated for 24 h at 37 °C at 800 rpm after which oxidized products were analyzed quantitatively to determine the total degree of oxidation in the LPMO-treated chitin. The rests of the 20 g/L reactions were centrifuged in an Eppendorf centrifuge (13,000 rpm for 3 min), the supernatant was removed and the soluble products in the supernatant were subjected to degradation with 2 µM *Sm*ChiA and 0.5 µM *Sm*CHB as described above, to determine the amount of solubilized oxidized products. The pelleted oxidized chitin was washed with buffer (3 × 1 mL of 50 mM sodium phosphate pH 7.0) by repetitively suspending the chitin in buffer and removing the supernatant after centrifugation. Finally, the oxidized chitin was suspended in buffer to 20 g/L. Again, samples were taken from all six reactions and diluted in buffer supplemented with 2 µM *Sm*ChiA and 0.5 µM *Sm*CHB to a substrate concentration of 2 g/L. The reactions were incubated for 24 h at 37 °C at 800 rpm and the resulting samples were used to determine the amount of insoluble oxidized products). Compounds in *Sm*ChiA/*Sm*CBH degraded samples was quantified as described below.

### Quantitative analysis of GlcNAcGlcNAc1A

Prior to product quantification, LPMO-generated products, were degraded with only *Sm*CHB (soluble fractions) or a combination with *Sm*ChiA and *Sm*CHB (for total or insoluble fractions) to yield the oxidized dimer (GlcNAcGlcNAc1A) and non-oxidized monomer (GlcNAc). Analysis and quantification of chitobionic acid (GlcNAcGlcNAc1A) were carried out using an RSLC system (Dionex) equipped with a 100 × 7.8 mm Rezex RFQ-Fast Acid H+ (8%) (Phenomenex, Torrance, CA, USA) column operated at 85 °C. Samples of 8 µL were injected to the column and sugars were eluted isocratically using 5 mM sulphuric acid as mobile phase with a flow rate of 1 mL/min. Standards of GlcNAcGlcNAc1A (10-500 μM) were used for quantification. GlcNAcGlcNAc1A was generated in-house by complete oxidation of *N*-acetyl-chitobiose (Megazyme; 95% purity) by the *Fusarium graminearum* chitooligosaccharide oxidase (*Fg*ChitO) as previously described (47, 48).

### Determination of apparent melting temperature (Tm)

The apparent melting temperature (T_m_) of the proteins was determined according to a protein thermal shift assay (ThermoFisher Scientific) based on using SYPRO orange, a fluorescent dye, to monitor protein unfolding (49). The quantum yield of the dye is significantly increased upon binding to hydrophobic regions of the protein that become accessible as the protein unfolds. The fluorescence emission (RFU) was monitored using a StepOnePlus real-time PCR machine (ThermoFisher Scientific). T_m_ was calculated as the temperature corresponding to the minimum value of the derivative plot (–d(*RFU*)/dT vs T; Figure S2). 0.1 g/L LPMO in 50 mM sodium phosphate buffer (pH 7.0) was heated in the presence of the dye in a 96-well plate from 25 °C to 95 °C, over 50 minutes. For each protein, the experiment was carried out in quadruplicates (i.e. n = 4).

### Binding studies with CjCBM5 and CjCBM73

Binding studies were performed as previously described (20). The equilibrium binding constants (*K*_d_) and binding capacity (B_max_) were determined for *Cj*CBM5 and *Cj*CBM73 by mixing protein solutions of varying concentrations (0, 20, 50, 75, 100, 150, 300 and 500 µg/ml for CjCBM5 and 0, 10, 20, 50, 75, 100, 150, and 300 µg/ml for *Cj*CBM73; protein concentration determined by A_280_) with 10 mg/ml pre-oxidized (see above) or untreated α-chitin. Before adding the chitin, A_280_ was measured for each of the prepared protein solutions (in 50 mM sodium phosphate buffer, pH 7.0), to create individual standard curves for each protein. After addition of chitin, the solutions were placed at 22 °C in an Eppendorf Comfort Thermomixer set to 800 rpm for 60 min. Subsequently, samples were filtered using a 96-well filter plate (Millipore), and the concentration of free protein in the supernatant was determined by measuring A_280_. All assays were performed in triplicate and with blanks (buffer and 10 mg/ml α-chitin). The equilibrium dissociation constants, *K*_d_ (µM), and substrate binding capacities, B_max_ (µmol/g α-chitin), were determined by fitting the binding isotherms to the one-site binding equation where P represents protein: [*P*_*bound*_] = *B*_*max*_[*P*_*free*_]/(*K*_*d*_ + [*P*_*free*_]), by nonlinear regression using the Prism 6 software (GraphPad, La Jolla, CA).

### NMR spectroscopy

NMR spectra of 70 µM *Cj*CBM5 and *Cj*CBM73 in NMR buffer (25 mM sodium phosphate and 10 mM NaCl, pH 5.5) containing 10 % D_2_O were recorded at 25 °C on a Bruker Ascend 800 MHz spectrometer with an Avance III HD (Bruker Biospin) console equipped with a 5 mm Z-gradient CP-TCI (H/C/N) cryogenic probe at the NV-NMR-Centre/Norwegian NMR Platform at NTNU, the Norwegian University of Science and Technology, Trondheim, Norway. ^1^H chemical shifts were referenced internally to the water signal, while ^13^C and ^15^N chemical shifts were referenced indirectly to water based on the absolute frequency ratios (50). Backbone and side chain assignments of *Cj*CBM5 and *Cj*CBM73 were obtained using ^15^N-HSQC, ^13^C-HSQC, HNCA, HN(CO)CA, HNCO, HN(CA)CO, CBCANHHNCACB, CBCA(CO)NH and H(C)CH-TOCSY.

For *Cj*CBM5 the BEST (band-selective excitation short-transient) (51) versions of HNCA, HN(CO)CA, HNCO, HN(CA)CO and HN(CO)CACB were recorded. The assignments have been deposited in the BioMagnetic Resonance Databank (BMRB) under the IDs 34519 (*Cj*CBM5) and 34520 (*Cj*CBM73).

### Structure elucidation

The NMR data were recorded and processed with TopSpin version 3.6 and analyzed with CARA version 1.5.5 (52). For structure determination, 3D ^13^C-edited and ^15^N-edited NOESY-HSQC spectra, as well as 2D ^1^H-^1^H NOESY spectra were recorded. NOE cross-peaks were manually identified, assigned, and integrated using the NEASY program within CARA version 1.5.5. Dihedral torsion angles (φ and ψ) were calculated from chemical shift data (C^α^, C^β^, H^N^, H^α^, H^β^, N and C’) by TALOS-N (53). Structures were calculated using the torsion angle dynamics program CYANA 3.97 (54). The structure calculation started by generating 200 conformers with random torsion angles, and the dihedral angles in each conformer were optimized using simulated annealing in 10,000 steps, to fit the restraints. The 20 conformers with the lowest CYANA target function values were energy-minimized using YASARA (55), first *in vacuo*, followed by using water as the explicit solvent and calculating electrostatics by applying the particle mesh Ewald method (56). In both these steps the YASARA force field (57) was applied. The coordinates of the minimized CBM conformers have been deposited in the Protein Data Bank (PDB) under the IDs 6Z40 (*Cj*CBM5) and 6Z41 (*Cj*CBM73). The two structures were aligned using the combinatorial extension (CE) algorithm, which determines the longest continuous alignment between fragment pairs (58).

### Titration of CBMs with chitohexaose

The interaction between *Cj*CBM5 and chitohexaose, (GlcNAc)_6_, was investigated using NMR spectroscopy. A ^15^N-HSQC spectrum was recorded of a sample of ^15^N-labeled *Cj*CBM5 (70 µM) in 50 mM sodium phosphate containing 10 % D_2_O and used as reference. Another sample of ^15^N-labeled *Cj*CBM5 (70 µM) with 10 mM (GlcNAc)_6_ in 50 mM sodium phosphate containing 10 % D_2_O was prepared. After recording a ^15^N-HSQC spectrum of this latter sample, it was mixed with the reference sample to obtain the following concentrations of (GlcNAc)_6_ 0.2, 0.5, 1.0 and 5.0 mM, while maintaining a constant protein concentration. A new ^15^N-HSQC spectrum was recorded at each (GlcNAc)_6_. The chemical shift perturbations (Δδ) were calculated using the equation: 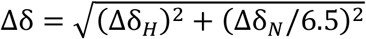, where Δδ_*H*_ is the change in chemical shift for the amide proton and Δδ_*N*_ for the amide nitrogen, in ppm (59). The *K*_*d*_ was estimated using Gnuplot 5.2 (www.gnuplot.info) based on an average of the amide chemical shift perturbation (Δδ) from the five most affected amino acids (W283, T284, Q285, Y296 and G297). The function used for fitting was Δδ = Δδ_*max*_[*S*]/(*K*_*d*_ + [*S*]), where Δδ_*max*_ describes the binding capacity as the maximum value of Δδ. Error bars in the chemical shift measurements correspond to 0.003 ppm.

The same procedure was applied for the NMR titration of *Cj*CBM73 with (GlcNAc)_6_, using *Cj*CBM73 (500 µM) and the following (GlcNAc)_6_ concentrations: 0.2, 0.6, 1.2, 2.8, 5.6 and 6.5 mM.

### Modelling coarse-grained α-chitin

The crystal structure of α-chitin at 300 K (60) was used to generate an all-atom α-chitin surface composed of 12 chains with 20 residues each, in UCSF Chimera version 1.13.1 (61). The all-atom model was coarse-grained using the bead mapping and topology parameters proposed by Yu & Lau (62) and bead types from the coarse-grained Martini version 3.0.beta.4.17 force-field (29). A rectangular simulation box was defined with the same size as the surface in the y (110 Å) and z (100 Å) dimensions and 150 Å in the x dimension.

### Coarse-grained simulations

The coarse-grained Martini version 3.0.beta.4.17 force-field was used in combination with GROMACS 5.1.4 (63) to simulate interactions between the CBMs and α-chitin. The constructs were coarse-grained using the Martinize2 program (64). An elastic network model (65) was used to constrain the overall structure of the CBMs. The beads in the chitin surface were kept in place by applying a harmonic potential with a force constant of 1000 kJ mol^-1^ nm^-1^ on the x, y and z positions. Models of *Cj*CBM5 or *Cj*CBM73 were manually placed in the simulation box above the chitin surface by using PyMOL (66). The simulation box was filled with water beads and the system was neutralized with beads corresponding to Na^+^ and Cl^-^ ions to an ionic strength of 0.15 M. The complex was energy-minimized using a steepest-descent algorithm (100 steps, 0.03 nm max step size) prior to being relaxed for 1 ns, with a time step of 5 fs, using the velocity-rescale (v-rescale) thermostat (67), Parrinello-Rahman barostat (68) and Verlet cutoff scheme (69). Simulations were run on the relaxed models with a time step of 20 fs, using the v-rescale thermostat, Parrinello-Rahman barostat and Verlet cutoff scheme in the NPT ensemble. Frames were written every 1 ns for each trajectory. Representative conformations of coarse-grained models of *Cj*CBM5 and *Cj*CBM73 were backmapped to atomistic models by using the CG2AT2 program (70).

### Adjusting the protein-chitin interaction strength in the Martini force-field

Initially, binding between CBMs and chitin was not observed, therefore, drawing inspiration from Larsen et al (71), we considered the following approach to modify the Martini version 3.0.beta.4.17 force-field to promote CBM binding to chitin. First, the interaction strength (i.e. ε parameter in the Lennard-Jones potential) between chitin beads and protein beads was increased by 10% and well-tempered metadynamics simulations (WT-MetaD; see below) were run until binding between the CBMs and chitin was observed. Then, a binding path for each CBM, which included bound and unbound conformations was selected from the WT-MetaD simulations, and umbrella-sampling (US) simulations were performed on these conformations as described below. Finally, we tested different interaction strengths by generating topologies where the interaction strength was modified from 0% (unchanged) to 15% increase of the chitin-protein interaction strength and ran umbrella-sampling simulations for each interaction strength. Figure S3 shows the free-energy surfaces calculated for each umbrellas sampling simulation using the weighted histogram analysis method (WHAM) (72, 73). Dissociation constants calculated from each free-energy surface are shown in Table 1.

### Well-tempered metadynamics (WT-MetaD) simulations

WT-MetaD simulations (74) were performed using the PLUMED 2.5 plugin (75–77) and the Martini model where chitin-protein interactions had been increased by 10%. We used the distance (*r*_*chitin*_) and number of contacts (cut-off distance, *r*_0_ = 0.7 nm) between aromatic residues in the putative substrate binding surfaces (*Cj*CBM5: Y282, W283, Y296; *Cj*CBM73: W371, Y378, W386) and the chitin surface in 15 µs long (15000 frames) simulations as collective variables to sample the binding of CBMs to chitin. Gaussian hills were added every 10 ps, with a starting height of 2.0 kJ mol^-1^, width of 0.5, and bias factor of 50. Figure S4 shows the evolution of the CVs and deposition of Gaussian hills over the course of the simulation.

Since the modeled chitin surface corresponds to the x,y-plane in the simulation box, *r*_*chitin*_, was calculated using Equation 1 (shown below with *Cj*CBM5 as an example), which uses the geometric center of the z-coordinates, *z*_*gc*_, calculated using Equation 2. Here, *z*_*i*_ is the z-coordinate of each amino acid or chitin bead, and *N* is the number of beads in the amino acid or chitin.

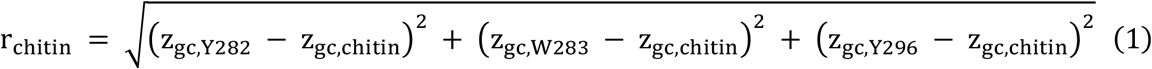

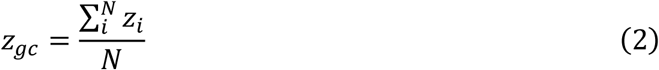

The number of contacts between all beads in each amino acid and all beads in the chitin surface, *c*_*i*_, were calculated using the COORDINATION routine in PLUMED 2.5 (i.e. Equation 3), with a cut-off distance *r*_0_ = 0.3 nm, *r*_*i*_ is the distance between all beads in each amino acid in the CBMs and all beads in the chitin surface.

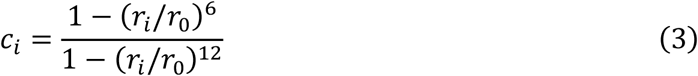

Weights for each frame, *w_i_*, were calculated from the bias in the WT-MetaD simulation using the REWEIGHT_BIAS routine in PLUMED 2.5 (see (78) and https://www.plumed.org/doc-v2.5/user-doc/html/_r_e_w_e_i_g_h_tb_i_a_s.html for details). The weighted average number of contacts, ⟨*c*⟩_*w*_, was calculated using Equation 4 and errors in ⟨*c*⟩_*w*_ were estimated using block analysis (30).

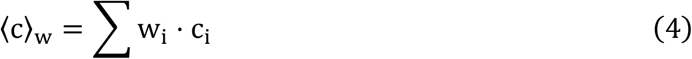

### Dissociation constants from umbrella-sampling simulations

A binding path for each CBM that included bound and unbound conformations was selected from the WT-MetaD simulations. Umbrella-sampling simulations were performed by running twenty-three 100 ns-long replicas using the PLUMED 2.5 plugin, where the distance between the putative binding surface and the chitin surface (*r*) was restrained from 0.0 to 4.4 nm in steps of 0.2 nm using a harmonic restraint with a force constant of 100 kJ mol^-1^ nm^-1^. Free-energy surfaces were calculated using the weighted-histogram analysis method (WHAM) (72, 73), where the errors were estimated by Monte-Carlo resampling. Dissociation constants (*K*_*d*_) were calculated from the free-energy surfaces (Figure S3) by using Equations 5-7, where *r*_0_ = 0·0 nm (i.e. the minimum distance that for which WHAM calculated a non-zero probability), *r*_*c*_ = 1·85 nm, *k*_*B*_ is Boltzmann’s constant, T is the temperature in Kelvin, *P*_0_ is the protein concentration in the simulations, N_A_ is Avogadro’s number, V_box_is the volume of the simulation box, and 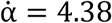 nm is the upper limit of *r*chitin.

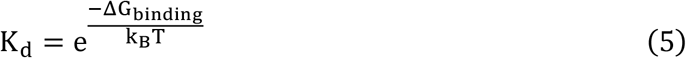

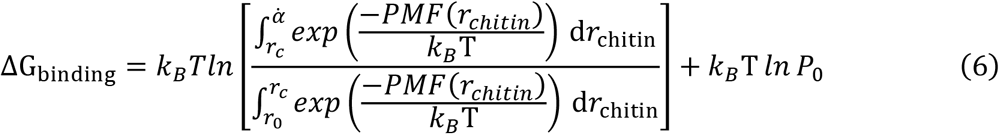

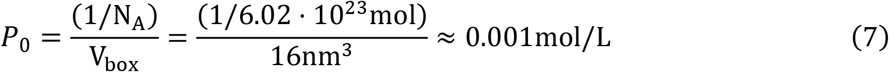

All the data and PLUMED input files required to reproduce the simulation results reported in this article are available online at https://github.com/gcourtade/papers/tree/master/2021/CBM5-CBM73-MetaD-US and on PLUMED-NEST (www.plumed-nest.org), the public repository of the PLUMED consortium (75) as plumID: 21.015.

## Acknowledgments

This work was funded by The Novo Nordisk Foundation, project numbers NNF18OC0055736 (ZF) and NNF18OC0032242 (GC) and by the Research Council of Norway through projects 226244, 247001, 262853 & 269408. Y.W. and K.L.-L. were supported by the BRAINSTRUC structural biology initiative from the Lundbeck Foundation.

