## Supporting Information for "NMR structures and functional roles of two related chitin-binding domains of a lytic polysaccharide monooxygenase from *Cellvibrio japonicus*"

**Table S1** Restraints and structural statistics for the 20 best conformers of the NMR solution structure of *Cj*CBM5 (PDB ID: 6Z40) and *Cj*CBM73 (PDB ID: 6Z41)

| | *Cj*CBM5 | *Cj*CBM73 |
|---|---|---|
| **Total number of NOE distance constraints** | **435** | **656** |
| Intraresidue (\|i-j\|=0) | 192 | 365 |
| Sequential (\|i-j\|=1) | 168 | 114 |
| Medium-range (1<\|i-j\|<5Å) | 25 | 61 |
| Long-range (\|i-j\|≥5Å) | 50 | 116 |
| **Torsion angle restraints** | **83** | **66** |
| Structure statistics (20 conformers) | | |
| CYANA target function value (Å$^2$) | 3.40 ± 0.36 | 2.83 ± 0.18 |
| Maximum residual distance constraint violation (Å) | 0.45 | 0.28 |
| Maximum torsion angle constraint violation ($^o$) | 5.75 | 4.21 |
| PROCHECK-NMR Ramachandran plot analysis [a] | | |
| Residues in favored regions (%) | 70.0 | 76.6 |
| Residues in additionally allowed regions (%) | 26.3 | 22.3 |
| Residues in generously allowed regions (%) | 1.1 | 1.1 |
| Residues in forbidden regions (%) | 1.6 | 0.0 |
| RMSD to the average coordinates (Å) | | |
| N, $C^{\alpha}$, C’ [a] | 1.47 ± 0.27 | 1.12 ± 0.28 |
| Heavy atoms [a] | 2.00 ± 0.29 | 1.52 ± 0.39 |
| N, $C^{\alpha}$, C’ (secondary structure)[b] | 0.45 ± 0.15 | 0.36 ± 0.11 |
| Heavy atoms (secondary structure)[b] | 1.35 ± 0.20 | 0.74 ± 0.13 |

[a] Residues *Cj*CBM5: 251-307; *Cj*CBM73: 338-397

[b] Residues *Cj*CBM5: 271-274, 276-280, 298-301 and *Cj*CBM73: 357-359, 362-366, 371-374, 390-394

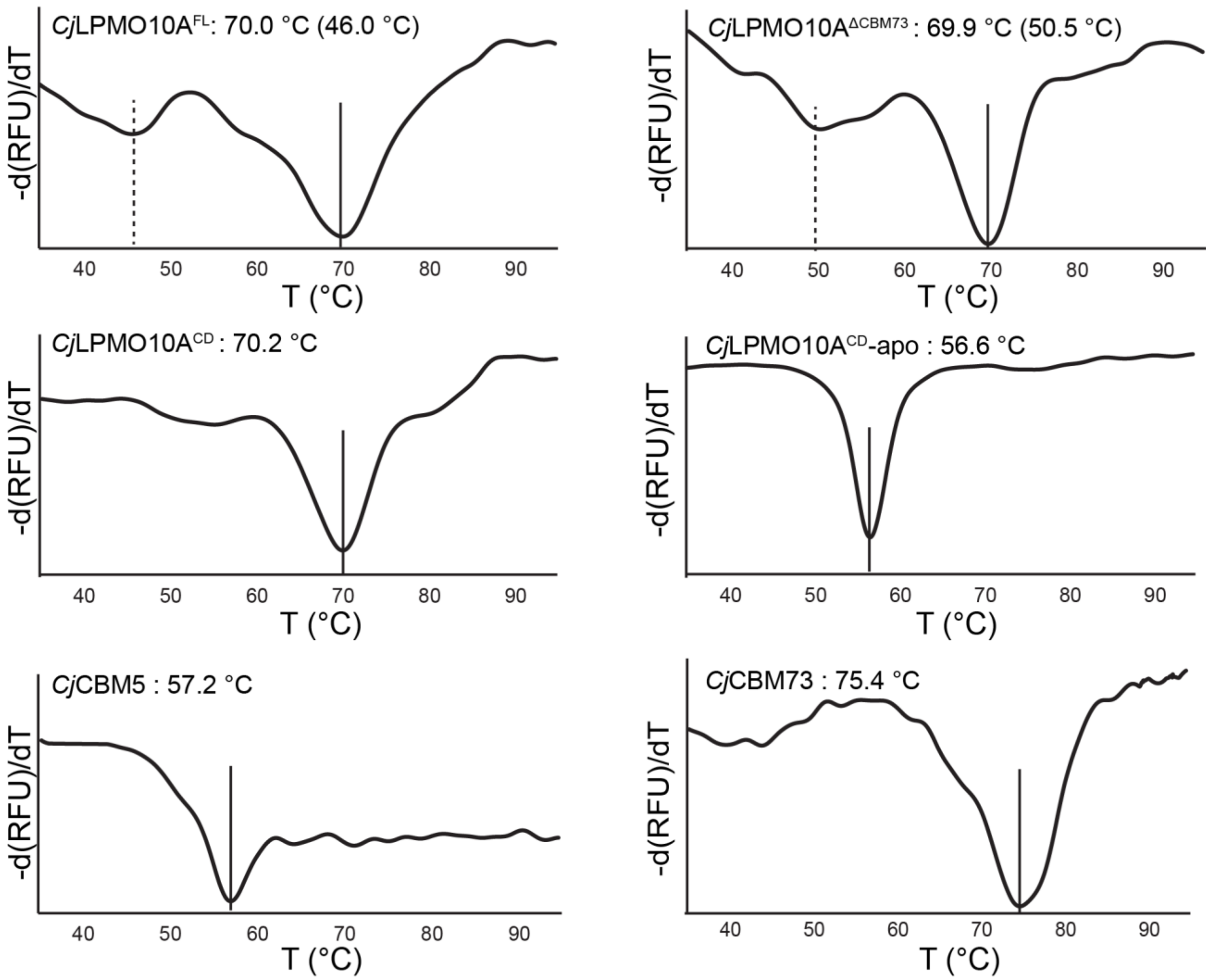


**Figure S1.** Thermal stability of CjLPMO10A variants. The plots show the melting curves and apparent melting temperatures ($T_m$) for copper-saturated *Cj*LPMO10A$^{FL}$, *Cj*LPMO10A$^{\Delta CBM73}$, and *Cj*LPMO10A$^{CD}$, as well as the apo form of *Cj*LPMO10A$^{CD}$ and non-metallated *Cj*CBM5 and *Cj*CBM73. The derivative of the fluorescence signal (-dRFU/dT, where "RFU" stands for relative fluorescence units") is plotted as a function of the temperature (1). The reactions contained 0.1 g/L protein and were heated from 25 °C to 95 °C, at a rate of 1.5 °C /min, in the presence of SYPRO orange (a fluorescent dye). The scans were performed four times for each protein and the Figures show a typical scan. All apparent melting temperatures ($T_m$) had standard deviations below ±0.3 °C.

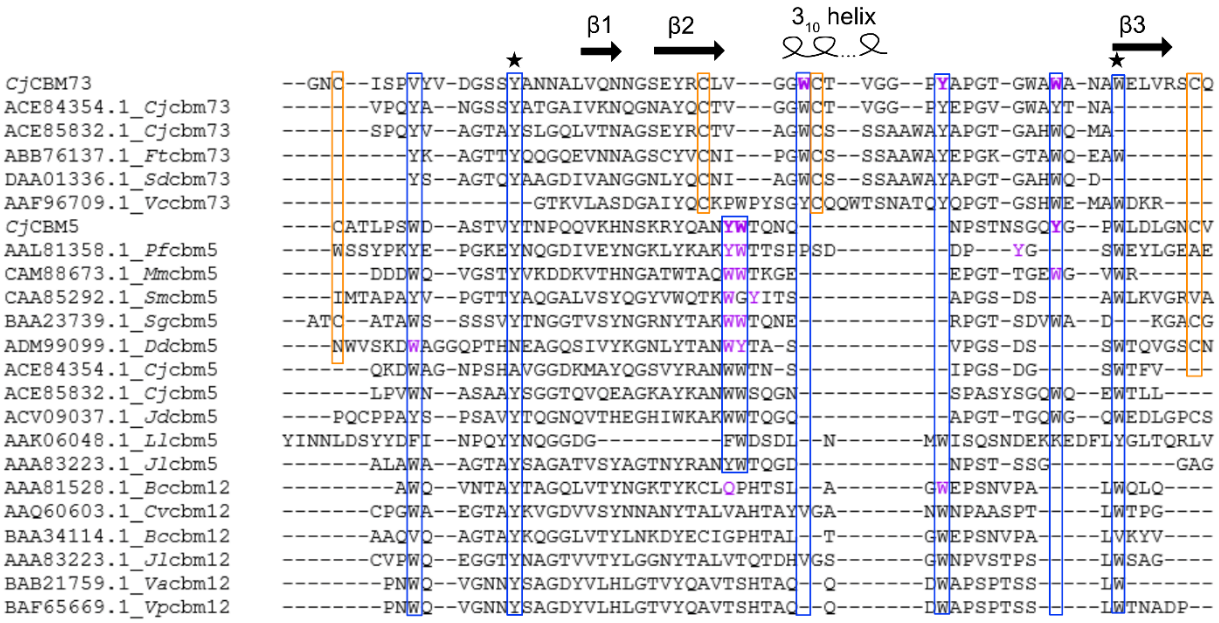


**Figure S2.** Multiple sequence alignment performed on a selection of members of CBM5, CBM12 and CBM73. The sequences are identified by the Genbank accession code and the CBM family registered in the CAZy database (2). The source organisms are abbreviated as *Cellvibrio japonicus* (*Cj*), *Francisella tularensis* (*Ft*), *Saccharophagus degradans* (*Sd*), *Vibrio cholerae* (*Vc*), *Pyrococcus furiosus* (*Pf*), *Mortella marina* (*Mm*), *Serratia marcescens* (*Sm*), *Streptomyces griseus* (*Sg*), *Dickeya dadantii* (*Dd*), *Jonesia denitrificans* (*Jd*), *Lactococcus lactis* (*Ll*), *Janthinobacterium lividum* (*Jl*), *Bacillus circulans* (*Bc*), *Chromobacterium violaceum* (*Cv*), *Vibrio alginolyticus* (*Va*), *Vibrio parahaemolyticus* (*Vp*). Aamino acids on the binding surface are highlighted in purple and correspond to the residues highlighted in Figure 8. These and other conserved aromatic residues are marked with blue boxes. The asterisks indicate two aromatic residues highly conserved in these CBMs, all buried in the hydrophobic core of these proteins. Conserved cysteines are marked with orange boxes. The MSA was prepared using Clustal Omega 1.2.4 (3).

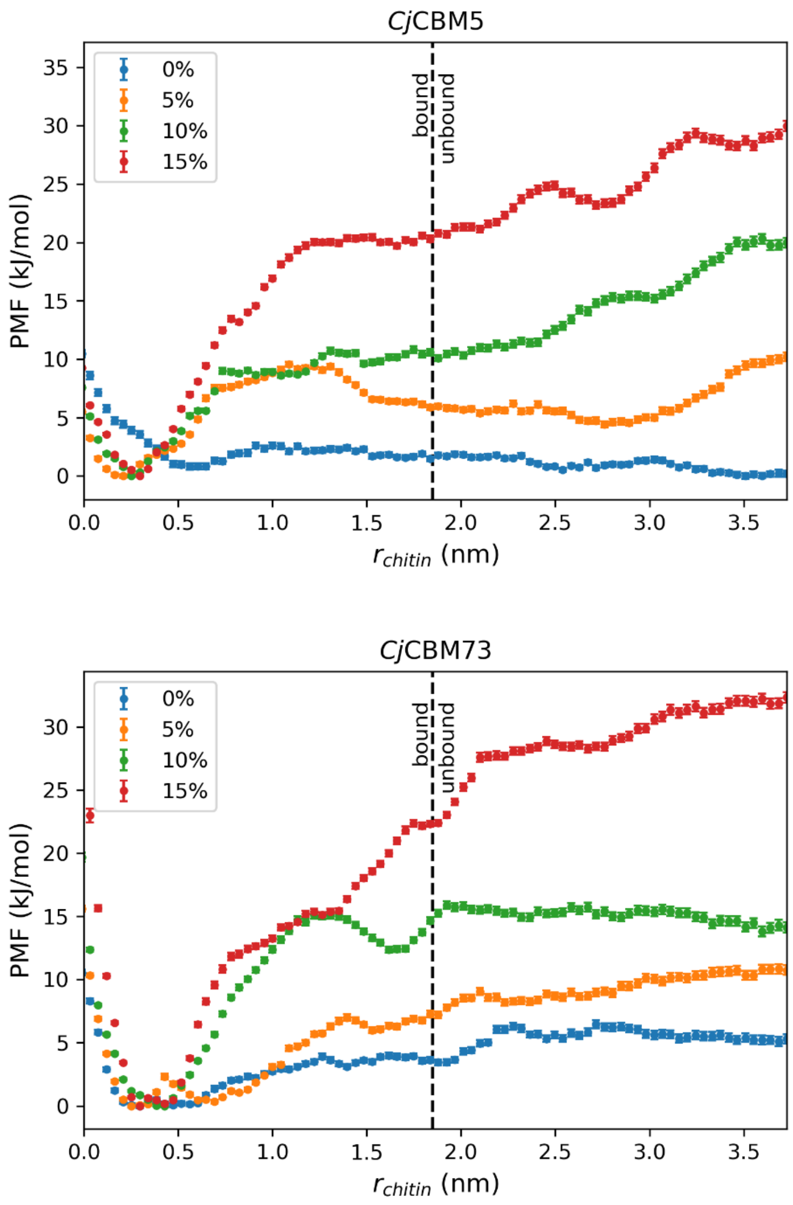


**Figure S3.** Potential of mean force (PMF) profiles for *Cj*CBM5 (top) and *Cj*CBM73 (bottom) as a function of their distance to α-chitin. The free-energy surfaces were calculated from umbrella sampling simulations performed at different chitin-protein interaction strengths: 0% (unchanged), 5% increase, 10% increase and 15% increase, and analyzed using the weighted-histogram analysis method (4, 5). The collective variable ($r_{\text{chitin}}$) is the Euclidean distance between the z-coordinate of the geometric center of the beads belonging to aromatic amino acids on the putative binding surfaces (*Cj*CBM5: Y282, W283, Y296; *Cj*CBM73: W371, Y378, W386) and the *z*-coordinate of the geometric center of all chitin beads. The cut-off value for between the bound and unbound states, $r_c$ = 1.85 nm, is indicated with a dashed line.

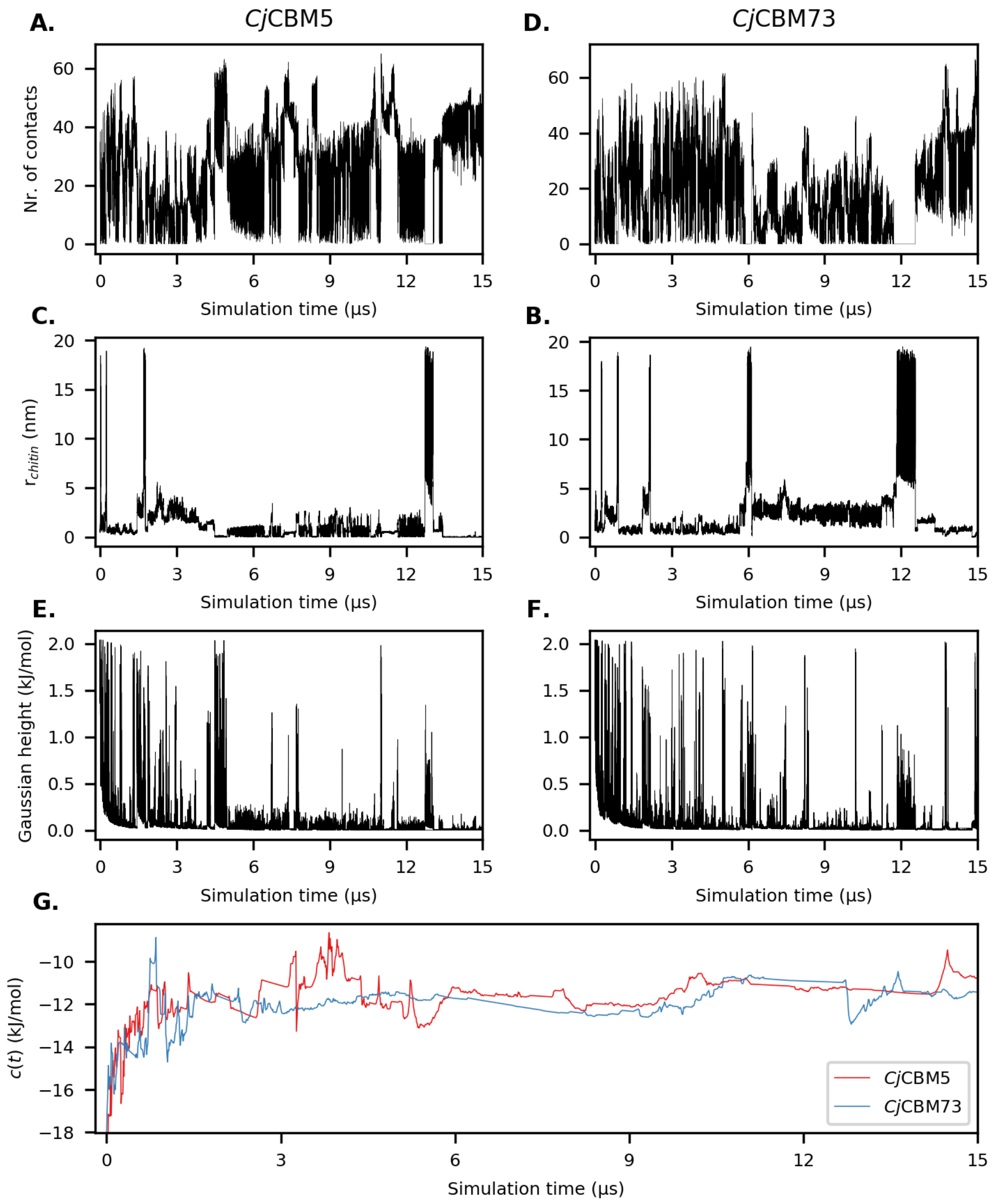


**Figure S4.** Time series of the number of contacts between chitin and selected amino acids on the surface of (A) *Cj*CBM5 (Y282, W283, Y296) and (B) *Cj*CBM73 (W371, Y378, W386), (C,D) the distance between these amino acids and the chitin surface ($r_{chitin}$), and (E,F) Gaussian hill deposition over the course of the WT-MetaD simulations. Panel G shows c(t), an estimate of the reversible work performed on the system at time t (6), calculated using https://github.com/lud0/comp-bio-tools/blob/master/reweight.py. The small number of transitions observed between the bound and unbound states of the CBMs, indicates that these simulations are not converged after 15 μs. However, for CjCBM5, both experiments and simulations find amino acids in the same region of the protein to be involved in chitin contacts (Figure 7).
